# SAMMBA is a high-throughput pipeline for isolating and phenotyping macroalgal strains

**DOI:** 10.1101/2025.09.04.674200

**Authors:** Cicero Alves-Lima, Luis Barreto, Carina Mónico, Lidiane Gouvêa, Francisca Felix, Brigitta Varga, Joana Filipe, Rita Camacho, Myrsini Lymperaki, Filipe Alberto, Leonardo R. Rörig, Aschwin H. Engelen, Ester A. Serrão, Gareth A. Pearson, Neusa Martins

**Affiliations:** Centro de Ciências do Mar do Algarve (CCMAR/CIMAR LA), Campus de Gambelas, Universidade do Algarve, 8005-139 Faro, Portugal; Department of Biological Sciences, University of Wisconsin-Milwaukee, Milwaukee, WI, United States; Laboratory of Phycology, Federal University of Santa Catarina (LAFIC – UFSC), Florianópolis, SC, 88.040-900, Brazil

**Keywords:** Seaweed Phenomics, Chlorophyll fluorescence, Breeding Automation, Marine biotechnology, Bioimaging

## Abstract

Anthropogenic climate change is causing the decline of seaweed forests in many parts of the world. Despite successful preservation efforts, their immense biodiversity is still severely underrepresented in germplasm biobanks throughout the world. These culture libraries can preserve genetic diversity and provide inoculum for marine forest restoration and mariculture ventures, and potentially accelerate the selection and breeding of climate-resilient and high-yielding strains. However, the complex life cycles and body plans of seaweeds pose a huge challenge for the development of standardized phenotyping and isolating protocols for microscopic stages, especially with the efficiency necessary to deal with the current pace of global climatic changes. Here, we present SAMMBA (Seaweed Automatable Microplate Microscopy for Breeding Approaches), an end-to-end pipeline for the high-throughput isolation, phenotyping and storage of macroalgal cells in 384-well plates (384WP). By optimizing fluorescence microscopy imaging and analysis, along with a novel fragmentation method and dilution-to-extinction isolation, different unialgal seaweed tissues could be regrown after thousand-fold dilutions. In a single plate, we successfully isolated 68 singlet gametophyte fragments of *Laminaria ochroleuca* (39 males, 29 females; 17.7% efficiency) and 60 spores of *Phyllariopsis purpurascens* (31.25% efficiency). Furthermore, the taxonomic versatility of SAMMBA was demonstrated through the successful isolation of 60 unialgal cultures of red algae (*Halymenia sp., Hydrolithon sp., Erythrotrichia sp.*) and 10 strains of the green alga *Ulva sp,* without cross-contamination. The viability and unialgal nature of the isolated strains were verified by distributing a single *L. ochroleuca* strain across an entire 384-well plate and imaging each well over 30 days. We found that the average specific daily growth rates (daily SGR) per well were 0.130 ± 0.006 and 0.117 ± 0.01 day^-1^ for males and females, respectively, showing a significant difference between sexes (n = 768; p = 1.27e^-53^), while edge effects significantly reduced daily SGR in males but not in females. This approach dramatically increases experimental reproducibility and statistical power compared to conventional methods. Due to its modular design and cost-effectiveness, SAMMBA is readily adaptable to macroalgal repositories globally. It supports high-throughput, selective recovery of unialgal strains without reliance on robotic platforms, while remaining fully compatible with automation. This system significantly expands the experimental and operational capacity in macroalgal hatcheries, providing a scalable foundation for phenomics, domestication programs, and standardized, verifiable biobanking of unialgal strains. Ultimately, SAMMBA could provide critical support for breeding strategies required to ensure the resilience of marine forests and aquaculture in a rapidly changing ocean.

## 1. Introduction

Anthropogenic activities are threatening global biodiversity at unprecedented rates, with projections estimating a 76-96% decline in marine macrophyte populations in temperate seas (1), particularly those with a high potential for tropicalization, where most marine forests are concentrated (2). In this alarming scenario, integrated strategies and policies have become essential to mitigate the impacts, preserve and restore these vulnerable marine socio-ecological systems (3). Among these, macroalgal germplasm biobanking represents an urgent and indispensable strategy to safeguard coastal marine biodiversity (4). It also plays a pivotal role in enabling the selective breeding of climate-resilient strains, which are essential for reforesting areas where biodiversity once flourished (5). Despite their recognized role as ecosystem engineers (6,7), macroalgae remain severely underrepresented in global *ex situ* collections. To support broader restoration and domestication efforts, it is crucial to expand germplasm banking efforts while ensuring greater taxonomic and functional diversity (8). This is especially important given that the nursery phase remains the most resource-intensive and technically demanding stage of any restoration program (9). The complex life cycles of many macroalgae, particularly the triggers of gametogenesis and sporogenesis in Rhodophyta and Ulvophyceae (10), further underscore the need for wider access to diverse, well-characterized strains to support both applied and basic research.

Developing fully standardized manipulation protocols across all macroalgal groups is likely unfeasible due to their vast phylogenetic, morphological and developmental diversity (11), although a number of widely adopted methodologies have emerged. These are primarily focused on kelps (Laminariales), *Ulva*, or commercially important red algae such as *Asparagopsis, Chondrus, Gracilaria, Palmaria* and *Porphyra* (12–16), likely driven by their relatively simple culturing requirements, high spore-release efficiency, and the easily identifiable microscopic propagules (spores, gametophytes, gametes and juvenile sporophytes). These characteristics facilitate reproducibility and scalability in experimental and applied contexts. Kelp gametophytes, for example, are typically settled onto glass slides following spore dehiscence on humid paper in darkness, and subsequently maintained in Petri dishes, where they can be manually monitored, isolated, and phenotyped under bright field microscopy (17). Smaller petri dishes (5cm) and multi-well plates (6-12 wells) have been used to downscale experiments, allowing multiple replicates and treatments (18,19). For *ex situ* collections, 5-100 ml sterile assay tubes have been used for long-term preservation, which increases stability of abiotic conditions and requires fewer media refreshment cycles per year (13,16).

More recent approaches have introduced high-throughput phenotyping (HTP) methodologies, such as heat-stress-tolerance (HST) evaluation by measuring chlorophyll autofluorescence (CAF) dynamics in 96-well plates (20), or spore isolation via flow cytometry (21). A similar HTP approach, integrating CAF and oxygen evolution, has recently enabled large-scale HST quantification in seaweeds such as *Ulva* and *Sargassum* (22). However, these emerging techniques often rely on specialized and costly instrumentation, limiting their accessibility and widespread use. In contrast, a custom and low-cost HTP method has been developed for some *Ulva* species, integrating phenotyping by imaging and metabolome approaches, marking another significant advancement for the macroalgal research community (23).

To advance these efforts and reduce the technical bottlenecks in strain isolation and characterization, we present a high-throughput, relatively low-cost workflow tailored for the nursery phase of macroalgal strains. This method integrates microtiter well plates (384 wells), chlorophyll autofluorescence microscopy and machine-learning-assisted image segmentation, by using the LabKit plugin (24) embedded in FIJI software (25). Together, these components enable a rapid and reliable isolation of multiple viable, unialgal strains in significantly less time than conventional approaches. In addition, it allows for precise monitoring of the unialgal nature of each isolate, and simultaneous phenotyping by growth measurements. By addressing key limitations in current isolation and phenotyping protocols, namely throughput, reproducibility, and accessibility, SAMMBA enables broad applications in strain isolation, storage and phenotyping, facilitating breeding and biodiversity conservation in macroalgal research.

## 2. Results

We present SAMMBA (Seaweed Automatable Microplate Microscopy for Breeding Approaches), a low-cost, end-to-end workflow for high-throughput isolation and phenotyping of macroalgal strains (Fig. 1). This approach integrates all essential components for isolating multiple macroalgal tissue samples using 384-well plates (384WP) as culture vessels, combined with automated image processing for quality control. Designed to enable manual microscopy screening, SAMMBA does not require motorized microscope stages or scientific-grade cameras, while images are processed entirely using open-source software. However, when automated microscopy with a motorised stage was used, the overall plate screening time was reduced by approximately two-thirds (data not shown).

**Figure 1.**
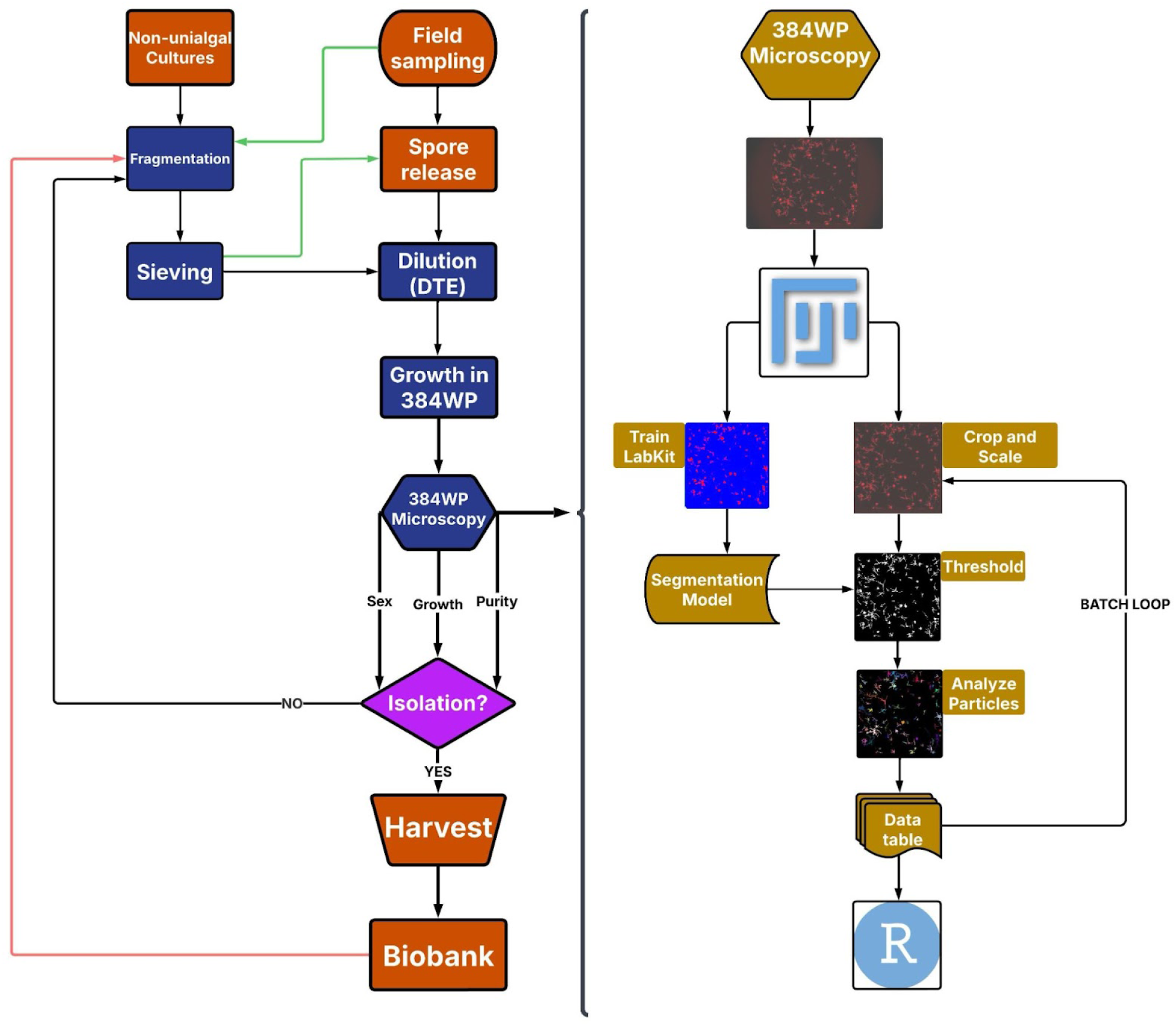
Flowchart of SAMMBA pipeline. . On the left, the sequence of steps is shown for isolating unialgal macroalgal propagules, starting either from field-collected or non-unialgal cultures, or for directly phenotyping isolated strains (red arrow). The green arrow represents the procedure for gametogenesis induction with *Ulva* thalli. Each blue box corresponds to a specific protocol, all of which are provided in the supplementary material. Red-colored boxes represent protocols that are not optimized in the SAMMBA framework; however, valuable recommendations are provided for these steps. Prior to dilution, the macroscopic tissue must be fragmented and sieved. The dilution step is always necessary to ensure reproducibility and avoid empty or overcrowded wells. A dilution-to-extinction (DTE) approach is specifically used for isolation to maximize the chances of finding unialgal propagules. The 384-well plate procedure is central to the entire procedure, allowing for the simultaneous monitoring of unialgal status and phenotyping by growth rate and developmental stages. On the right side, the image processing pipeline in FIJI is detailed, highlighting the integration of machine-learning model training by the LabKit plugin, as indicated by brown boxes. This pipeline is also provided as a FIJI macro script.

We first evaluated edge-effects on low volume 384WP wells caused by seawater evaporation, and salinity increase, as seaweed cultivation requires long term incubation. We then optimized image acquisition by fluorescence microscopy and processing workflows to allow a fast and reliable detection. Using this microscopy setup, we measured fragmentation yields of a microtube bead mill and compared to standard methods, optimizing a method for the disruption of multiple seaweed samples simultaneously. For the isolation, the fragmented seaweed tissue was subjected to a dilution-to-extinction (DTE) method to maximize the chances of finding unialgal propagules (singlet wells). The DTE method was then validated with *Phyllariopsis* spores, and red and green macroalgal microscopic propagules, showing a broad taxonomic applicability. We further validate the monitoring of all procedures by measuring growth rate across a full plate for up to 40 days, estimating edge-effects, purity, viability, and differences between sexes, strains and species with precision. We anticipate that these methods will enhance the quality of macroalgal physiology experiments, accelerating the development and establishment of new strains.

### 384-well plate edge-effects and Imaging

In order to quantify localized evaporation dynamics in a 384WP and its potential edge effects on seaweed viability, we measured volume decrease based on a methylene blue standard curve (Supp. Fig1-I). For efficient isolation, plates were covered with silicone seal mats (SSMs) and volume reduction was compared against uncovered wells. Volume reduction in peripheral wells followed an exponential trend during the first two weeks, as evidenced by markedly steeper regression slopes compared to the more gradual decline observed in central wells (Supp. Fig1A-F). Edge effects, measured by daily specific evaporation rates (SER), were evident in uncovered plates (Supp. Fig1B), and were significantly reduced by the use of SSMs (Supp. Fig1 C-F; Supp. Table 1). Highest SERs reached 3.3 % day^-1^ in a few peripheral wells with 80 µl (*mat80;* Supp. Fig1D), however global SERs were lower than the uncovered mat (*nomat100;* Supp. Fig1G). Notably, when analyzing central wells, *mat80* daily SERs were significantly higher than *mat50* and *nomat100*. On the other hand, central wells from *mat50* displayed the lowest daily SERs of all treatments and regions, which was confirmed by final salinities that didn’t differ significantly to the control, in both regions (Supp. Fig1H; Supp. Table 2).

We could only significantly reduce edge effects by using SSMs on *deep well* 384WP (Supp. Fig1H), as all edge wells had higher final salinity when compared to central wells (Supp. Table 2). However, when comparing the same regions from different plates, SSMs were efficient in reducing daily SERs and final salinity differences after 31 days. For convenience, unsealed 384WPs filled up to 100 µl were chosen for phenotyping macroalgal strains by short-term growth kinetics, eliminating the need to remove SSMs before microscopy. SSMs-plates with 50 µl had the lowest daily SERs in both plate regions. Longer storage can be achieved in deep well 384WP, as final salinities didn’t differ between edge and center wells, even after 4 months (Supp. Fig1H; Supp. Table 2).

To allow a precise distribution of fragmented gametophytes along several wells and to reduce noise, we implemented washing (by centrifugation with artificial seawater and BSA) and sieving steps (20 and 40 µm) to remove cell debris and larger fragments that may clog pipette tips and result in greater well to well variation in total tissue fluorescence. Then we established the precise quantification of the gametophyte area with an automatable fluorescence microscopy imaging method.

For imaging, we used a standard CMOS full frame camera (CANON EOS RP) and by using PI (propidium iodide) filter, chlorophyll autofluorescence (CAF) could be imaged as bright red or pale red for alive and dead gametophytes, respectively (Fig2A). High contrast to background was achieved, facilitating the identification of live cells which was confirmed by overlaying regions of interest (ROI) detected from PI over bright field (BF) images (Fig2-BC). This allowed a short exposure time of 0.5s, leading to a total time for capturing a single image per well of 4.68s, including the stage movement to subsequent well and focus adjustment. This resulted in a 30 minute screening for a whole 384WP. Only fluorescent images were taken. The LabKit image segmentation FIJI plugin (25,26) was used to create an automated segmentation model of individual gametophytes (n = 39), presenting no significant difference (W = 764, p = 0.9762), and a strong correlation (R^2^ = 0.998, p < 0.0001; Fig2G) to manual quantification.

**Figure 2.**
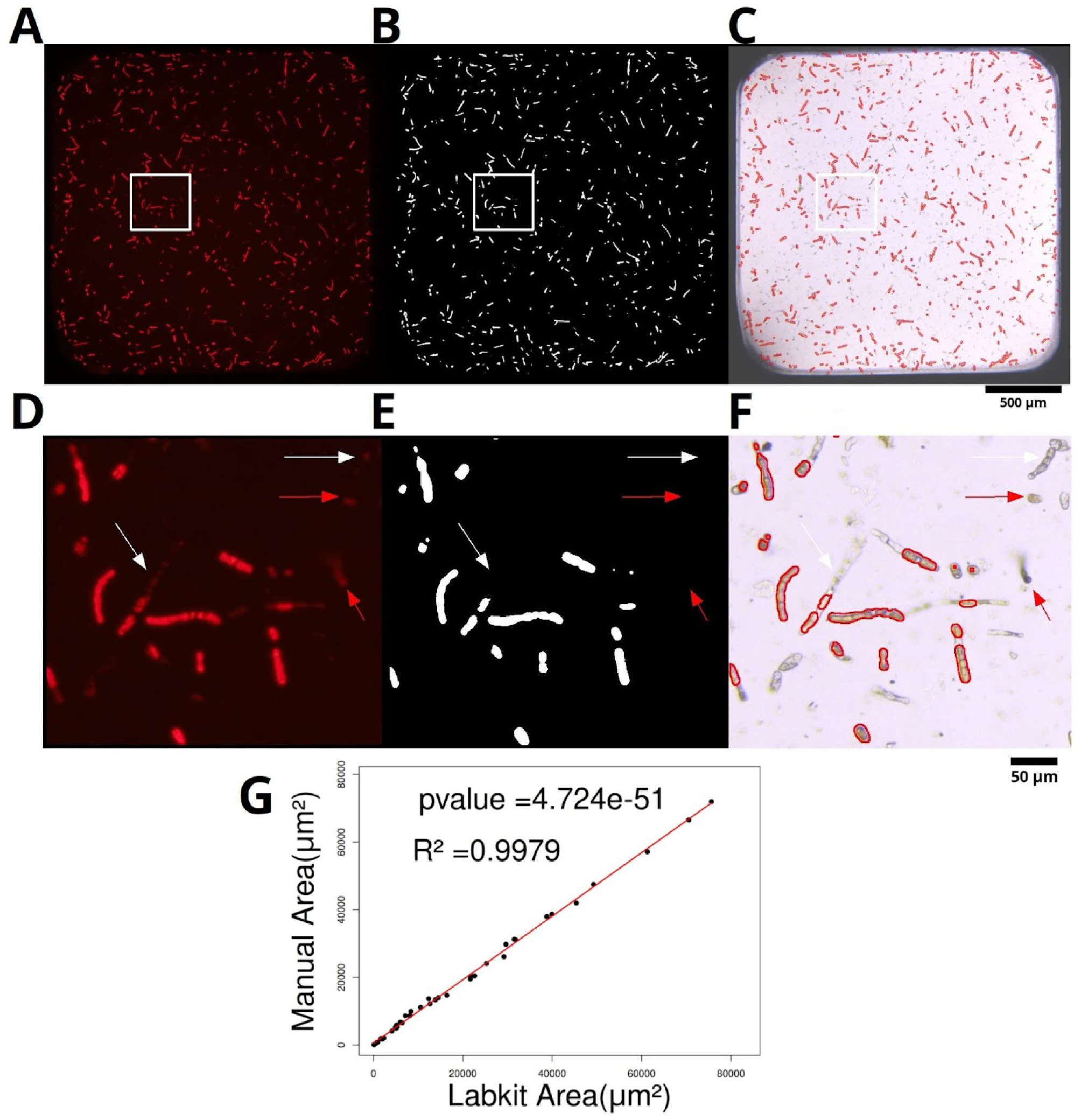
Validation of *Laminaria ochroleuca* gametophyte chlorophyll auto-fluorescence assay for high-throughput screening in 384 well plates. A, B and C are full-well views at 8x magnification (5x objective + 1.6x adapter) with white square indicating the region selected for the detailing in D-F, under 32x magnification. A) and D) show chlorophyll fluorescence images captured by the PI filter. In B and E, segmented binary image of each gametophyte after using the LabKit segmentation FIJI plugin. In C and F, the bright field microscopy image with the overlapped regions of interest (ROIs) derived from segmentation. Red arrows highlight examples of non-viable gametophytes that are pale red under PI filter that were successfully excluded from segmentation. Yellow arrows highlight dead gametophytes. In G, a strong linear regression (p-value = 4.72e^-51^; R^2^ = 0.9979) of manually segmented individual gametophytes and measured by LabKit plugin.

Although Bland–Altman analysis (Supp. Fig. 8) indicated bias with wide limits of agreement (–3556 to 2192.9), variance analysis revealed that only 7 of 39 measurements (>18%) differed by more than 10%. Importantly, the aim here is not to establish absolute agreement between methods, but rather to compare their ability to capture growth rates. In this context, what matters most is whether variation across time is proportional. Since the majority of measurements remained within acceptable variance, the relative changes over time are preserved, ensuring that growth rates derived from both methods remain consistent. This suggests that the observed differences have limited impact on the reliability of growth rate estimation is posterior analysis.

### Large-scale Fragmentation

To enable high-throughput processing of multiple *L. ochroleuca* gametophyte tufts, we optimized a large-scale fragmentation protocol and compared it to conventional methods. We evaluated the TissueLyser (QIAGEN), a bead mill designed for simultaneous disruption of multiple samples, typically used for seaweed biomolecule extraction (26,27). We explored frequency settings in order to maximize fragmentation efficiency while preserving maximum gametophyte viability. This method was compared to traditional protocols using mortar and pestle (MP) and a motorized grinder (MG).

Among all methods tested, fragmentation with mortar and pestle for 1 minute (MP1) was the most efficient, producing 4.1 times more fragments per well than the next best method: TissueLyser at 25 Hz for 3 minutes (TL25-3; Supp. Fig2A). However, the total recovered area did not differ significantly between MP1 and TL25-3, indicating that the latter did not cause tissue death by over-fragmentation of the tissue (Supp. Fig2B; Supp. Table 3). However, prolonged fragmentation (>1 min) led to significant losses in recovered area and counts in all methods, most probably as a result of cell lysis. Notably, the variation coefficient in fragment counts was around 20% for both MP1 (145.8 ± 25.4) and TL25-3 (35.5 ± 6.4).

### Dilution-to-extinction isolation

The initial density of the fragmented and sieved male and female gametophyte tissue was determined and submitted to a serial dilution-to-extinction (DTE) approach (Table1). The gametophyte dilutions followed a linear regression when compared to the logarithms of counts and area (Fig3A;D). High reproducibility up to 32x dilution subsequently declined, possibly related to non-homogeneous distribution (stochasticity) and/or differential adhesion to plastic surfaces (plate and tip) causing non-reproducible pipetting with highly diluted gametophytes.

**Table 1.**
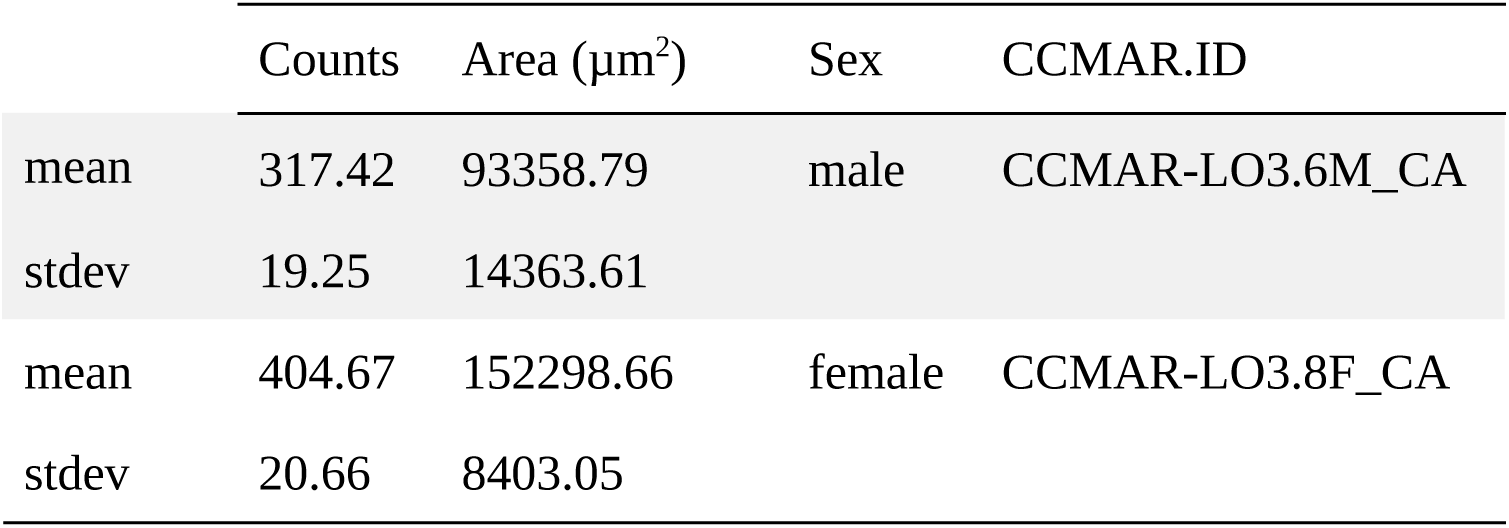
Inoculum densities in gametophyte counts per well (100ul) and total area (µm^2^) for both *Laminaria ochroleuca* strains used for the dilution-to-extinction isolation method (N = 12 for each strain).

**Figure 3.**
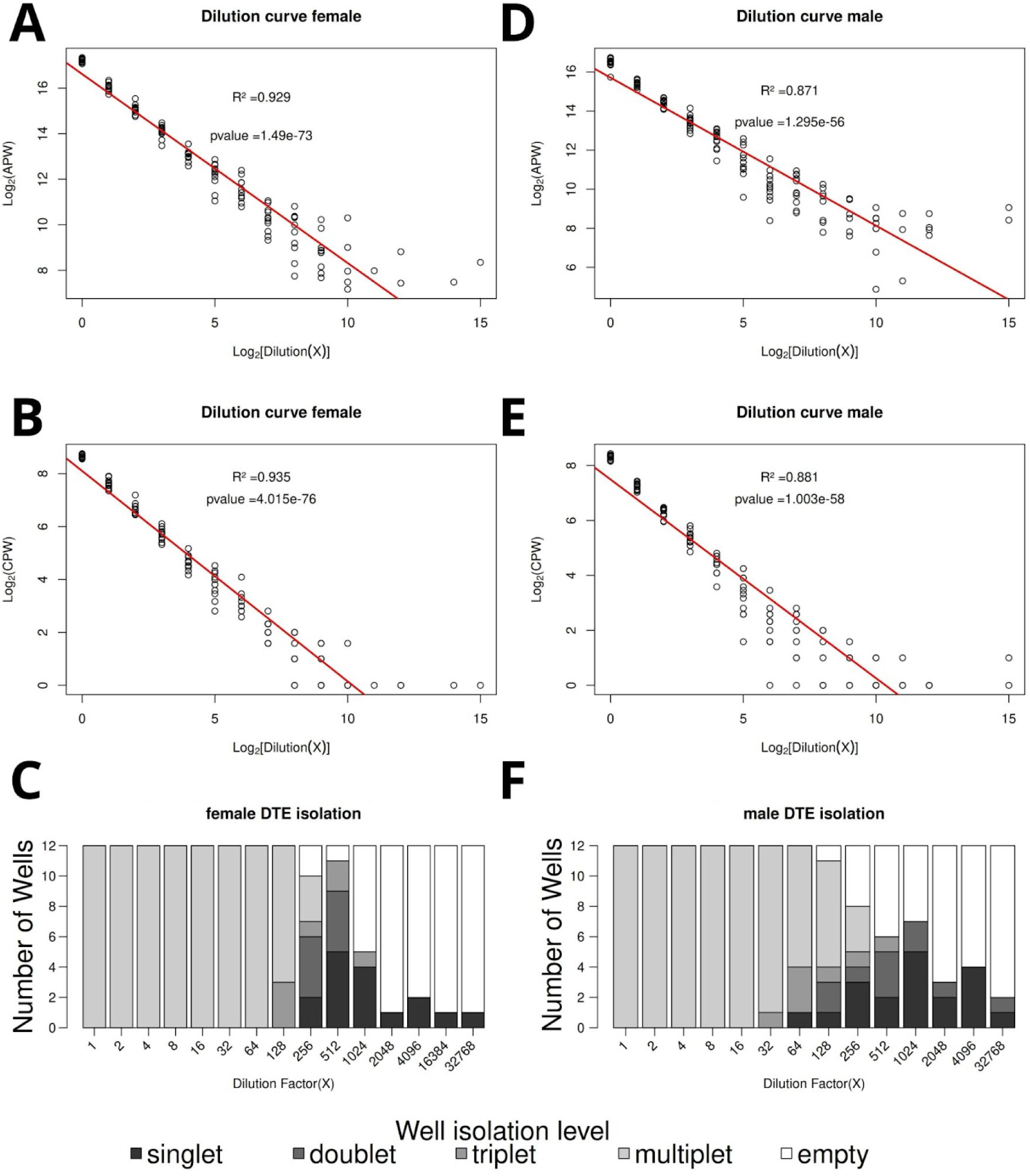
Isolation of *Laminaria ochroleuca* gametophytes by Dilution-to-Extinction (DTE) procedure. Female (A–C) and male (D–F) fragmented gametophytes are represented. Panels A and D show linear regressions of dilution against area per well (APW); B and E display counts per well (CPW). Bar plots in C and F illustrate the frequency of 384WP wells categorized by the counts of gametophytes: singlets, doublets, triplets, and multiplets (≥4). White bars represent empty wells. Based on inoculum densities (Table 1) and dilution levels yielding the highest singlet frequencies, optimal densities were defined as 7.89 (512x) and 3.49 (1024x) gametophytes per milliliter for females and males, respectively, to maximize singlet recovery.

We determined that the optimal dilution for maximizing the number of wells containing single gametophyte fragments (singlets) was ∼0.34 male gametophytes per well (equivalent to a 1024× dilution), and 0.78 female gametophytes per well (512x dilution). Considering that each well had 100 µl, the densities were 3.49 and 7.89 units/ml for males and females, respectively. Using these densities, fragmented tissues were distributed across half a 384-well plate (192 wells), yielding 39 male and 29 female singlets (Table 2). For the remaining 192 wells, we applied the 3.49 units/ml density to dilute and isolate *Phyllariopsis purpurascens* spores, assuming their distribution would more closely resemble that of *L. ochroleuca* male cells given their comparable size. This yielded 60 isolated spores, which subsequently developed into viable gametophytes (Table 2).

**Table 2.**
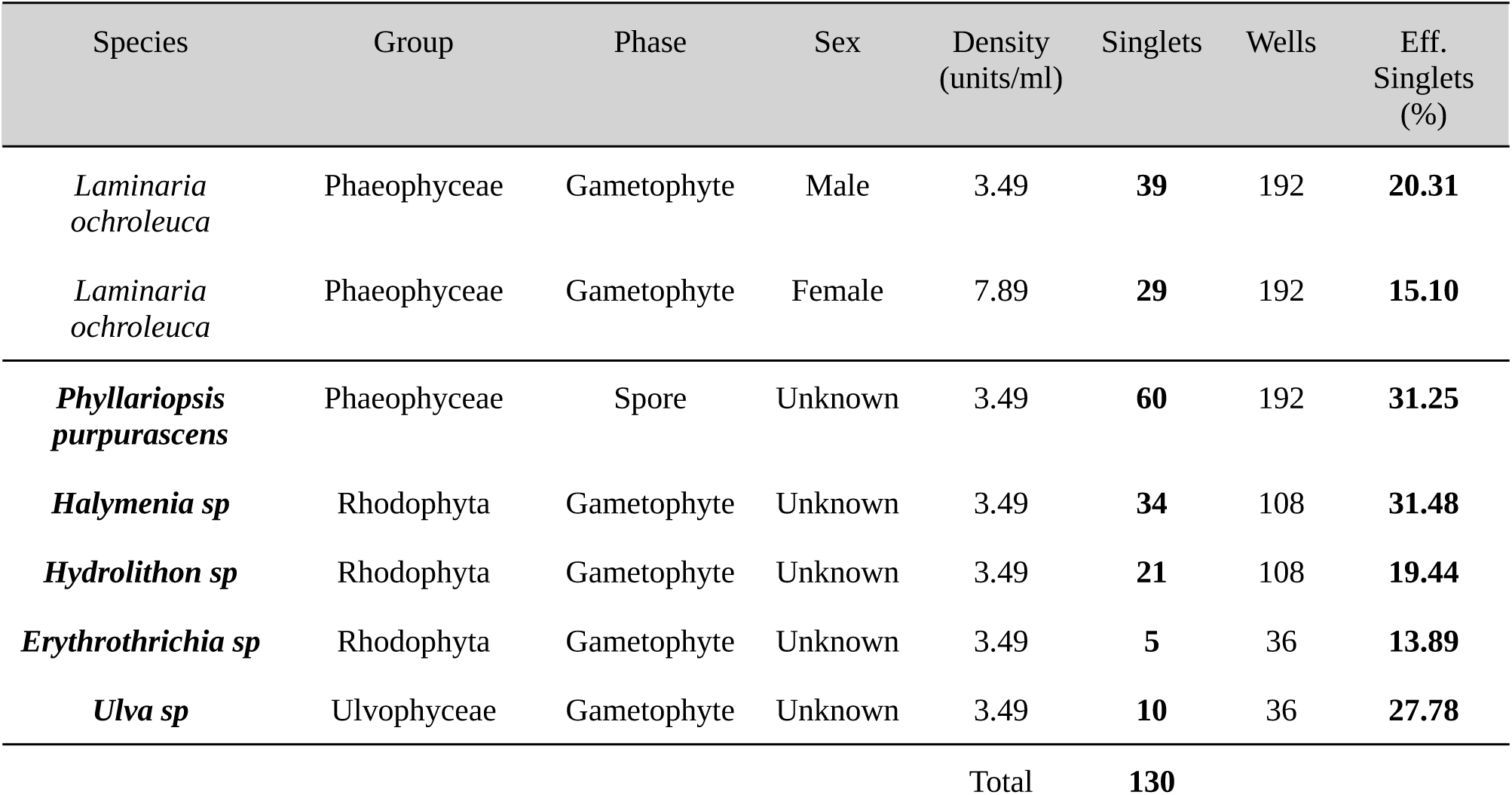
Overview of the isolation of all strains with the SAMMBA pipeline. The isolation efficiency (Eff. singlets) is the proportion of wells with only one seaweed fragment (singlet) on all wells used for the dilution-to-extinction. The densities (units/milliliter) were established for *L. ochroleuca* gametophytes and used for the other species highlighted in bold.

We also quantified the frequencies of wells containing two (doublets) and three (triplets) gametophytes, given that it is still feasible to manually isolate spatially separated fragments to increase isolation efficiency (Fig3C;F). Although the transfer of isolated gametophytes to larger culture flasks remains to be optimized, SAMMBA significantly outperforms traditional methods in isolation speed. Specifically, we achieved a transfer rate of 35 viable strains per hour using 384-well plates, compared to only 9 strains per hour when manually transferring from 8 cm Petri dishes, representing at least a fourfold increase in efficiency (data not shown).

The taxonomic versatility of SAMMBA was demonstrated through the successful isolation of 60 unialgal cultures of red algae (*Halymenia sp., Hydrolithon sp., Erythrotrichia sp.*) and 10 strains of the green alga *Ulva sp.,* also in a single plate (Fig. 4). All these isolates were monitored for 30 days and showed average daily SGRs of ca. 0.05 - 0.10 day^-1^ (Supp. Fig 6 and 7), highlighting the feasibility to phenotype a range of seaweed taxa by growth rate.

**Figure 4.**
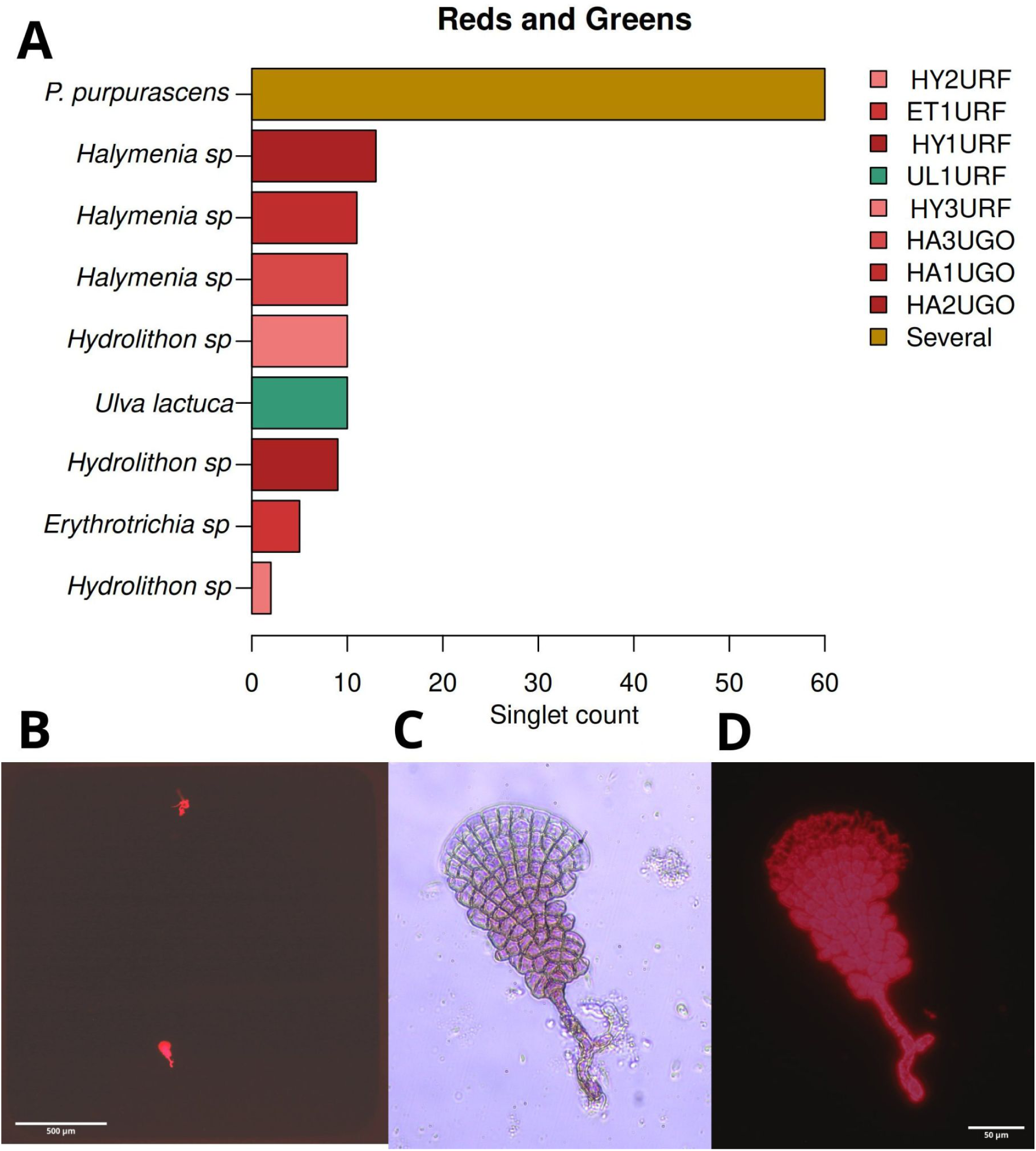
Validation of SAMMBA pipeline for the isolation of non Laminariales macroalgae. In A) the bar shows the number of singlet wells of 3 species of Rhodophyta, one *Ulva sp.* and the spores of *P. purpurascens.* The legend shows the CCMAR biobank coding for the isolated strains. In total, 130 potential new strains were added to the CCMAR biobank. In B, a full well view of a fluorescent doublet isolate from *Hydrolithon sp.* No other fluorescent algae can be seen in the well area. C, the lower isolate from B under bright field and 32x magnification. In C, the lower isolate from B at PI fluorescence under 32x magnification.

### High-throughput monitoring by growth rate

The low variation of the 32x dilution of the DTE curve allowed us to establish a protocol for the high-throughput evaluation of previously isolated *L. ochroleuca* gametophytes. At a density of 240 gametophytes/ml we could obtain high reproducibility along a whole plate pipetting, which proved to be a viable method for measuring growth rate of several samples in a single plate.

We could measure the area of the gametophytes for 30 days and plot growth curves to evaluate the precision of the LabKit model during seaweed development (Supp. Fig5). The plate plot shows a full growth curve for each well, constituted of all growth phases: the lag, exponential and stationary. Peripheral wells had earlier stationary phases, but the exponential phase slopes did not differ significantly from wells located in the center of the plate.

A high *intra-plate* reproducibility was found on the first day after fragmentation (DAF = 0) where total gametophyte counts per well (CPW), was 18.4 ± 0.27 and 20.3 ± 0.3 for females and males, respectively (mean ± SE; N=384). This was reflected in a high growth rate homogeneity across each 384WP plate, and revealed sex differences in growth rate. Although the average daily SGR differed only 11%, males grew significantly faster than females with daily 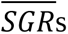 of 0.130 ± 0.006 and 0.117 ± 0.01, respectively (n = 768; p-value < 1.27e^-53^). For both groups, the growth rate showed to be continuous since the first week for all samples, certifying that a precise growth rate can be obtained within a week after fragmentation.

Early edge effects could be measured in the bottom row (P), which were represented by a stationary phase before DAF 21, followed by the top row (A). Wells in the 4 plate corners (A1, A24, P1 and P24) were also affected by earlier edge effects than others. Only male daily SGRs were significantly reduced by edge effects (Fig5E, Supp. Table 4). There was no correlation between daily SGR and gametophyte area on DAF 0, nor between daily SERs calculated from the evaporation optimization (Supp. Fig4). When growth rates were analyzed between smaller groups of wells, such as comparing all 24 columns (n=16; Fig5D), no significant differences were found between *inter-plate* groups while sex differences were still detectable, with exception of 2 groups (Fig5D; Supp. Table 5).

**Figure 5.**
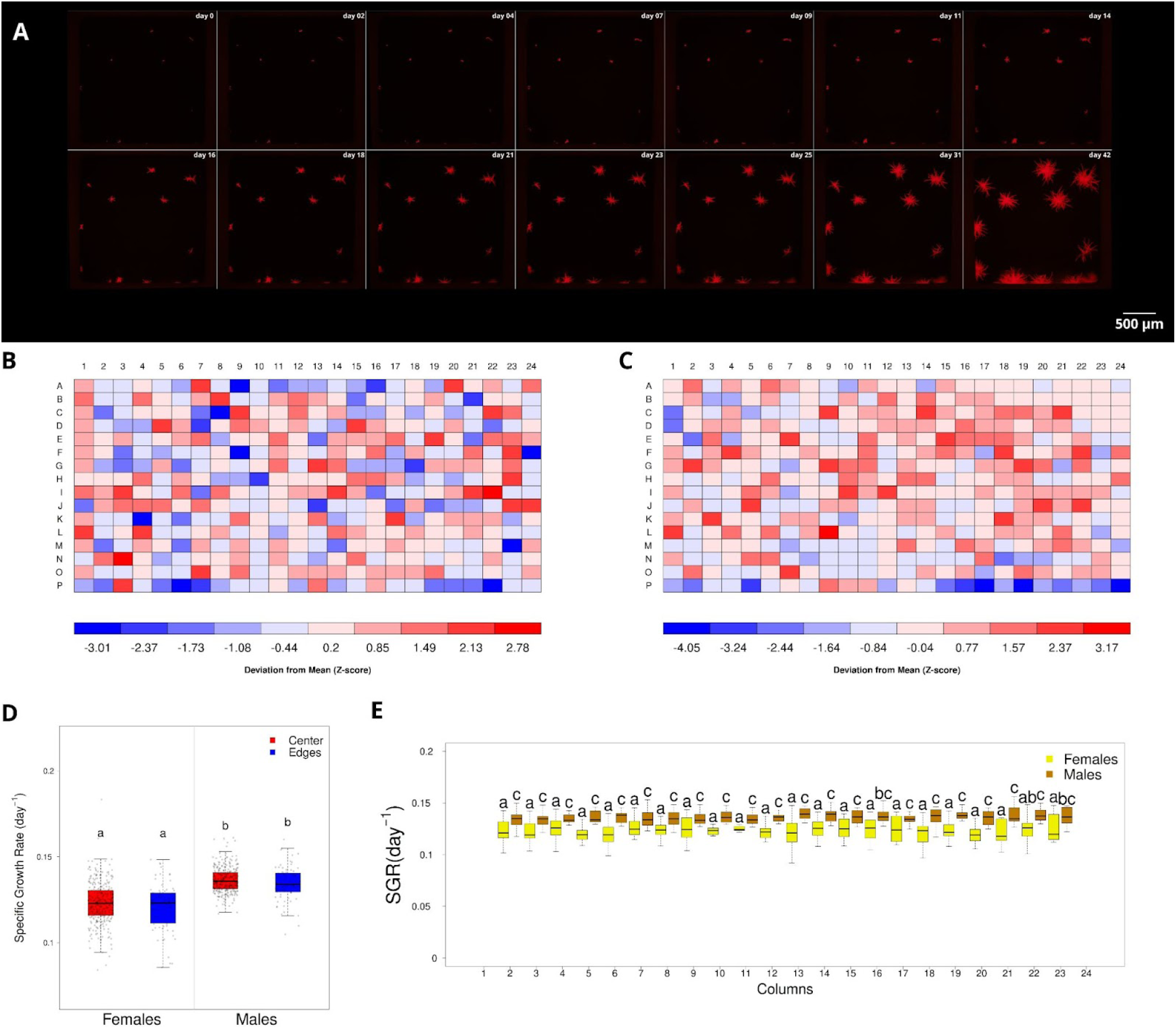
Growth rate analysis of *Laminaria ochroleuca* female and male gametophytes cultured in 384-well plates. (A) Frame montage illustrating the growth progression of a representative male gametophyte (panel C - K3) over 42 days. (B–C) *Plate-plot* showing spatial distribution of specific daily growth rates (daily SGR) for female (B) and male (C) gametophytes across the plate. Z-scores were computed for each well and visualized using red–blue palettes to reflect deviations from the mean daily SGR. The low Z-scores observed in most wells and the lack of a radial gradient indicate minimal edge effects. (D) Comparison of all daily SGRs between sexes reveals that males grew significantly faster than females (Wilcoxon test: W = 236.27, p-value < 1.27e^-53^) and wells located on plate edges had significantly lower growth due edge effects in males, but not in females. (E) Boxplots displaying the sex-biased differences in daily SGRs across plate columns, confirming maintenance of sex differences using only 16 replicates.

Growth rates were also monitored after fragmentation optimization and DTE isolation. MP1 fragmentation method showed the highest daily SGRs (0.148 ± 0.08), with mean values 15% greater than those observed for TL25-3 (0.127 ± 0.04), however no significant difference was found (Supp. Fig2C; p=0.06). The TL25-3 daily SGRs exhibited only 4.8% variation among all 16 replicates, the lowest of all tested conditions, highlighting its consistency and reliability. Following DTE isolation, *P. purpurascens* gametophytes exhibited a daily SGR of 0.054 day⁻^1^ (Fig. Supp. 6). For Rhodophytes and *Ulva* (Fig. Supp. 7), average SGRs were 0.08, 0.07, 0.11 and 0.07 day^-1^ for *Erythrotrichia, Halymenia sp, Hydrolithon sp* and *U. lactuca,* respectively. An isolate of *Hydrolithon sp* showed a maximum SGR of 0.22 day^-1^, the highest SGR recorded in this study.

## Discussion

Here we describe the optimization of SAMMBA, a comprehensive pipeline integrating diverse protocols for the isolation and phenotyping of macroalgal propagules and vegetative fragments in 384-well plates. SAMMBA establishes a novel high-throughput method for advanced germplasm biobanking, breeding strategies and fundamental biology. This method relies on the intrinsic clonality and totipotency of most macroalgae (16,28), enabling tissue disruption, dilution, and regeneration for reliable unialgal isolation. When applied to spores, the pipeline also enables scalable isolation of new strains. The pre-microscopy protocols are fully compatible with automation, reflecting SAMMBA’s flexibility and interoperability. This seamless integration across imaging platforms ensures that the system remains adaptable and scalable for diverse experimental needs.

The core advantage of SAMMBA lies in its imaging solution: the precise compatibility of the 384WP wells to a standard 35 mm full-frame sensor high resolution camera. By limiting seaweed growth to a minimum detectable space, this method provides a holistic and representative assessment of both fragment distribution and growth dynamics. This also avoids subsampling, random selection, and stitching routines required by smaller FOV cameras and large culture plates, enabling rapid, manual plate-wide screening which facilitates post processing. Full-well imaging ensures complete spatial coverage, improves the accuracy of quantification, and reduces both operator bias and variability introduced by field selection, enabling monitoring at the level of individual gametophytes/propagules.

We deliberately selected a commercial grade camera, which, while less sensitive than scientific models, reduces costs by more than 20-fold. When considering additional expenses of motorized stage and proprietary control software required by high-end setups, the SAMMBA approach becomes even more cost-effective. By prioritizing wide-field epifluorescence screening, we were still able to obtain a moderate signal-to-noise ratio due to sufficiently bright chlorophyll autofluorescence (Fig2A;D).

By capturing an entire 11.35 mm^2^ well (3.65 mm diameter) in a single frame, SAMMBA enables precise quantification of gametophyte density and its relationship to experimental treatments, an essential factor given the strong influence of cell spacing and sex ratio on fertility outcomes (29). This single-image resolution also allows individual propagules to be morphometrically analyzed, significantly increasing the accuracy and reproducibility of experiments. Furthermore, by aligning sequential images from growth kinetics assays, subtle cellular movements can be tracked, and high-quality time-lapse videos can be generated to visually validate phenotypic data. Taken together, these features advance seaweed phenotyping toward full compliance with FAIR (Findable, Accessible, Interoperable, and Reusable) data principles, facilitating data sharing and cross-laboratory validation, extending its impact beyond individual experiments (30).

### Imaging and segmentation

Imaging of chlorophyll autofluorescence was performed using a consumer-grade mirrorless camera coupled with standard inverted microscopy adapters and fluorescence filters, offering a practical and effective solution for high-throughput imaging. The system provided sufficient signal-to-noise ratio for reliable detection of CAF across 384WPs, with consistent exposure settings and controlled illumination. Although the use of a broadband filter set introduced some non-specific weak fluorescence of plate borders and micro fibers, it was effectively addressed during image segmentation through defining them as part of the “background” class.

In addition to being affordable and adaptable, this custom setup is suitable for large-scale screening applications where quantification of fluorescence area, rather than intensity, is the primary objective. While not indicated for precise photometric analysis, SAMMBA enables reproducible morphometry suitable for many applications. By using the PI filter, we tackled three main challenges of this technique: background autofluorescence and shading, gametophyte exposure to conditions that induce gametogenesis (31) and time to screen the whole plate. The use of an open-source platform such as FIJI for imaging from controlled laboratory phenotyping (32–34) to in situ field monitoring (35,36) also demonstrates the broad accessibility of SAMMBA.

LabKit provides a fast and accessible solution for image segmentation, enabling the detection and quantification of individual macroalgal tissues in fluorescence microscopy images. While recently developed CNN-based plugins such as DeepImageJ (37) offer advanced segmentation via deep learning, these typically require custom Python models, high-performance GPU hardware, and expertise in external machine learning workflows. Such dependencies limit accessibility and reproducibility in research environments without dedicated computational resources. In contrast, LabKit runs entirely within FIJI, offering a user-friendly, interactive interface that integrates seamlessly with standard image analysis pipelines and performs reliably on conventional desktop workstations. This makes it a broadly applicable and reproducible option for high-throughput segmentation tasks in macroalgal and other biological studies. However, LabKit’s pixel classification cannot reliably separate developmental morphotypes (males, females, oocytes, and sporophytes) under fluorescence as nearly identical red signals often appear mixed or overlapping within a single field of view (data not shown). Addressing this limitation will require the integration of deep learning approaches capable of resolving fine morphological differences, thereby enabling accurate quantification of fertility outcomes in controlled crosses.

Fluorescence microscopy also stands out as a straight-forward technique to precisely detect viable seaweed tissue. Most previous studies rely on bright field microscopy, where recently fragmented gametophytes can still appear pigmented, while being inviable.

### Fragmentation and Isolation

The throughput needed to take advantage of a 384WP system demanded a cumbersome procedure of manual fragmentation during the preparation of seaweed cell suspensions. Most seaweeds are multicellular and highly clonable organisms, therefore fragmentation has typically been used to generate experimental replicates by fragmentation (17,34). Mortar and pestles have been largely used, while the motorized portable pestle grinder was introduced more recently (13). The TissueLyser offers precise control over fragmentation intensity, and has been widely used for DNA/RNA extraction from disrupted tissue, but not for physiological experiments. We show here that the high throughput of the TissueLyser compensates for lower fragmentation performance: TL25-3 (25Hz during 3min) allows the processing of up to 16 samples per minute (48/3) using dual 24-well racks. Although MP1 (Mortar and Pestle during 1min) produced 4.1x more fragments, the overall processing speed of TL25-3 results in approximately 3.9x greater efficiency. The practical limitations of MP such as the need for autoclaving, cleaning, and storage of multiple MP sets further reduce its feasibility and scalability. Additionally, TL offers less manipulation as samples can be stored immediately without the need for transfer to another container.

Following controlled disruption, tissue regeneration was achieved via the isolation of sieved fragments through the DTE approach. Cycles of fragmentation, sieving, dilution and regeneration can be repeated multiple times on the same strain, ensuring isolation to the unialgal level (Fig1). Repetition of isolation protocols is a common procedure, for example the scraping, apex excision and regrowth is an effective method to isolate unialgal strains of Gracilarian seaweeds (38).

We were able to isolate tissue fragments and spores from 6 species of seaweed, two Phaeophyceae (*Laminaria ochroleuca* and *Phyllariopsis purpurascens)*, three Rhodophyta (*Erythrotrhichia*, *Hydrolyton* and *Halymenia*) and 1 Ulvophyceae (*Ulva sp).* All non-phaeophyceae seaweed were manipulated and grown under the optimized protocol for *L. ochroleuca* gametophytes.

The high degree of replication allowed by the SAMMBA system, enables parallel cultivation of multiple subcultures from the same strain, making it possible to identify and discard contaminated or aberrant wells before finishing isolation or without compromising the established strains from the collection. Purity assessments are highly reliable under these conditions, particularly when paired with imaging and specific staining protocols.

SAMMBA greatly facilitates the recovery, isolation, and cultivation of macroalgal strains by offering a streamlined and high-throughput format that simplifies previously laborious procedures. Strains that were often lost due to low viability or overgrowth by contaminants, can be effectively rescued and reestablished. The 384WP system enables isolation of single filaments or fragments into individual wells, dramatically improving the ability to purify target strains and, in the case of sexually dimorphic species like kelps, identify with precision the sex of each strain. For species historically considered challenging to culture, like *Phyllariopsis sp* (39), SAMMBA’s modular and scalable format offers testing of multiple medium variables like nitrogen, phosphate, iron and pH in the same plate. Also, several plates occupy just a tiny fraction of an incubator shelf, allowing multiple temperature and irradiance tests simultaneously.

SAMMBA enabled monitoring of both viability and purity for over 40 days by maximizing well volume under controlled humidity conditions, thereby minimizing evaporation. The use of zip-seal bags further reduces evaporation, allowing cultivation in standard incubators (without humidity control), which are typically more affordable. In addition to reducing water loss, higher culture volumes ensure greater nutrient availability and minimize meniscus-related edge shading during bright field microscopy, enhancing image quality and consistency (Fig2C).

The use of multiwell plates and dilution for isolation of unialgal, and even axenic, macroalgae is a standard procedure to manually isolate low density culture of gametophytes (16). In the SAMMBA framework, we adapted this approach by employing substantially lower initial densities, thereby increasing the likelihood of obtaining singlet wells. From a single thousand-fold dilution of spores or fragmented gametophytes, hundreds of replicate cultures can be established. The entire volume of the diluted inoculum can be distributed across all wells of a high-density microplate, increasing the chances of establishing unialgal cultures by a near-complete sampling.

### High-throughput Phenotyping

The successful isolation of algae requires continuous monitoring, as cryptic contaminants may overgrow early stages, compromising putatively isolated strains (40,41). Moreover, some unialgal strains fail to grow under standard conditions such as Provasoli medium (12), limiting downstream applications. To address these challenges, we optimized SAMMBA for frequent and rapid monitoring in 384-well plates. By applying precise dilutions, we established inoculation densities that ensured reproducible area-based growth measurements across wells. Importantly, seaweed microscopic tissues exhibit strong adhesion, mediated by sulfated polysaccharides (42), which can hinder reproducibility when pipetting gametophyte fragments. This issue was minimized by adding BSA during fragmentation, resulting in high reproducibility during subsequent DTE isolation.

Evaluating intra-plate variability by inoculating the same strain across a full plate could establish 384WP as a reliable platform for physiological experiments. By using 384 replicates, we precisely measured the influence of edge effects on seaweed growth and compared inter and intra-plate differences. Statistical differences in daily SGR between sexes were detected when the average diverged by as little as 11%. Column-wise comparisons with reduced replication (n = 16) also showed the same pattern. The results obtained conform with previous measures indicating higher growth in male versus female kelp, as well as higher resilience to thermal stress (43–46).

This comparative analysis of growth curves could reveal lower growth rates for *P. purpurascens* (Fig. Supp 6), which is likely attributable to the culture temperature (13 °C), which is close to the species’ typical lower thermal limit in temperate warm waters (39,47). However, more tests are needed in order to reveal detailed thermal tolerance for survival and development of this scarcely studied kelp.

High-throughput assays in 384WP are commonly used for screening various biological samples such as bacteria, yeast, human cells and fish (48–51). However, long-term growth assessment exposes macroalgae tissue to increased salinity due to evaporation in low culture volumes. Although salinity is not a significant stress on kelp sporophytes (52,53), gametophytes have only been assayed for hyposaline stress tolerance (54,55). In coastal marine ecosystems, hypersaline stress is rare but can impact seaweed physiology in intertidal pools (56) and in the vicinities of desalination plants discharges (57). We measured salinity up to 45 PSU in outer 384WP wells after 4 months without media renewal. Even using silicone seal mats, hypersaline conditions are a threat to gametophytes cells. We therefore provide guidelines and procedures to reduce evaporation and replenish media rapidly, allowing the manual processing of 4 plates per hour (i.e., 4 x 384 samples).

Relative growth rate is a common metric to evaluate stress tolerance in kelps, and to establish optimal gametogenesis conditions in sporophytes (58), but growth rate measurements are scarce at the gametophyte level (34,43,44,46). As area and length measurements are currently measured manually, optical aberrations (shading and vignetting) and contamination can pose serious challenges when determining gametophyte sex and viability, making accurate physiological comparisons difficult. An important way to overcome this limitation is by the use of chlorophyll autofluorescence, which has already allowed the screening of kelp strains directly in the gametophyte phase (59). SAMMBA is plate reader-compatible, allowing a wide comparison to these previous studies. Furthermore, by allowing the simultaneous test of many culture conditions, the workflow supports the optimization of growth media and environmental parameters in a dedicated way for each strain.

## 4. Limitations and Perspectives

The SAMMBA workflow presents some operational challenges, from plate preparation to image post-processing. Preventing cross-contamination and ensuring consistent volume handling require careful training to achieve reliable performance, particularly when frequent media renewal is necessary. However, when robotic pipetting arms are available, these limitations can be largely eliminated. For physiological experiments, minimizing media renewal is highly recommended to reduce labor and risks of contamination.

Evaporation is a persistent concern in 384WPs due to low culture volumes. For extended incubation beyond 30 days, particularly in deep-well plates, silicone sealing mats (SSMs) are crucial to mitigate seawater salinity increases. However, condensation often accumulates on the internal surface of the SSMs, increasing the risk of cross-contamination during media exchange or sampling. We showed here that a brief centrifugation before removing the mats resulted in viable seaweeds, and reduced droplet transfer between wells. Additionally, SSMs reduce irradiation by approximately 50%, a factor that must be considered when programming light settings in growth chambers. Failure to adjust for this light attenuation may exacerbate edge effects caused by unequal illumination and temperature gradients, which are further intensified under limited gas exchange.

Effective use of SAMMBA requires meticulous plate planning. While fully random distribution of replicates and controls is discouraged due to known asymmetrical edge effects (60), some level of randomness is highly recommended. This can be overcome by pre-pipetting the whole volume for all replicates of each strain in a 96 deep-well plate and planning the distribution.

Stage movement during microscopy may introduce focal plane variability, necessitating frequent Z-axis adjustments or the use of a calibrated focus map for automated navigation. We obtained an average manual imaging time per plate of ca. 30 min, which demands sustained attention or advanced automation to ensure image consistency. This was 65% slower than the automatic screening (20 min) using a motorized stage on another microscope. These timeframes require the control of room temperature to avoid thermal stress.

The image analysis component of SAMMBA can be computationally intensive. A single full plate can yield approximately 900 MB of fluorescence images per filter, each requiring up to 2 GB of RAM for segmentation when using LabKit. Therefore, for multi plate processing, access to a High-Performance Computing (HPC) cluster is advisable, especially as LabKit supports headless operation (24). Managing the large data volumes generated, particularly for time-series growth curves, also necessitates robust data storage and analysis workflows. We provide concise and detailed R! scripts to assist these steps.

Understanding these technical aspects underscores SAMMBA’s transformative potential. The platform enables the controlled screening of hundreds of replicates for early-stage phenotyping of growth, pigmentation, or morphology, accelerating the development of new seaweed strains. By combining established staining and labeling techniques using chromogenic substrates, fluorophores, and dye-conjugated probes (61), a wide array of features can be quantified, including cell wall polysaccharides, lipid vesicles, mitochondria, chloroplasts, viable cells, vacuoles, cytoskeleton dynamics and vesicle transport.

SAMMBA also supports fine-scale exposure of cultures to environmental gradients such as temperature, light, or nutrient availability. This allows researchers to characterize growth dynamics, stress resilience, and developmental transitions at a level of throughput previously unachievable in macroalgal systems. The platform is particularly suited to dose–response and time-course studies for pollutants such as pharmaceuticals, heavy metals, or microplastics, providing highly replicated, low-reagent experiments ideal for ecotoxicological risk assessments or regulatory frameworks.

By supporting the establishment of more diverse germplasm banks, SAMMBA is critical for ecological restoration initiative, as strains more tolerant to temperature, acidification and with higher adhesion to rocky substrate can be selected. Its broad taxonomic applicability can change the underrepresentation of other key macroalgal lineages such as Rhodophyta and Ulvophyceae in breeding programs. Together with key management tools for climate-smart marine restoration (62) it can provide higher chances of success in such complex activity. Given the rapid pace of environmental change, tools like SAMMBA are urgently needed to close the gap between current seaweed cultivar development and emerging ecological demands (43).

In addition, SAMMBA offers significant benefits for microbiome research. Miniaturized macroalgae-microbes co-culture assays can reveal specific interactions as hundreds of parallel combinations can be tested to identify growth enhancement, metabolite profile change, or influence on development. High-content imaging and fluorescence markers enable spatially resolved observations of microbial localization on or within macroalgal tissues, supporting investigations into host–microbe symbioses or stress responses. SAMMBA also allows the visualization of the interaction between macroalgae to non-photosynthetic microorganisms such as fungi, by the simultaneous capture of both bright field and fluorescence images.

One of the most compelling applications of SAMMBA lies in the potential overcoming the long-standing challenge of developing stable macroalgal genetic transformation protocols (19). We show here that the platform allows parallel testing of up to 20 transformation conditions with 12 replicates (excluding two outer well rings), which can be useful to testing different plasmids, delivery systems, and chemical treatments, thereby increasing the likelihood of identifying successful recombinant strains. Once transformation is achieved, SAMMBA can also be used to screen edited lines, supporting in-depth functional studies of gene regulation. Moreover, SAMMBA allows phenotyping of both haploid and early diploid sporophytes, helping to clarify the genotype–phenotype correlations across life cycle stages.

SAMMBA’s methodological orchestration extends far beyond isolation and phenotyping. As seaweed biotechnology, conservation, and climate adaptation strategies evolve, SAMMBA offers a scalable platform that can support both faster fundamental discoveries and translational applications in the blue bioeconomy. By enabling broad access to advanced tools, SAMMBA promotes the decentralization of HTP capacity, empowering a wider range of research groups and fostering collaborative and inclusive scientific community.

## 2. Materials and methods

### 384-well Plate optimization: Evaporation assay

As gametophyte cultivation in 384-well microplates (**384WP**) has not previously been tested to our knowledge, we first checked the viability of the method by quantifying and minimising edge effects. We quantified localized evaporative volume reduction per well in flat and clear bottom transparent 384WPs (VWR: 732-3736) covered with silicone seal mats (**SSM**; Axygen® AxyMats; PN AM-384-DW-SQ). This was assayed by changes in methylene blue (30μM) absorbance (665nm) diluted in Artificial Sea Water (**ASW**; Tropic Marin® Classic Sea Salt N° 10134; Lot 33452016) over 42 days. A standard curve (Supp. Fig1-I) with decreasing volumes (100 to 20 μl) was used as a reference to calculate the volume reduction.

As SSMs cap plugs project inside the wells, the maximum volume was reduced from 120 µl to 90 µl, therefore we also tested how evaporation would vary with different volumes, namely 50, 80 (with SSM) and 100 µl (without SSM). We also tested the viability of deep well 384 well plates (VWR: 732-3327) with SSM and filled to 100 μl for long term storage. The humidity was stabilized at 50% by isolating plates with lids with Parafilm® and keeping them in transparent plastic zip-seal bags as reported previously (49).

The plates were centrifuged for 2 min at 3700 RPM and the SSMs were removed and the lid placed back, before inserting in a microplate reader (Biotek Neo2). By the end of the experiment, the salinity in outer wells edge wells (A1, A24, P1, P24) and in central wells (H12, H13, I12, I13) was measured by pipetting 20 μl in a hand-held salinity refractometer (ATAGO®).

To quantify specific evaporation rates in 384-well plates, the time-dependent decline in well volume was modeled using nonlinear least squares (nls() in R!) with an exponential decay function:

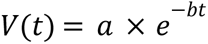

where V(t) is the remaining volume at time t (in days), a represents the estimated initial volume, and b is the specific volume decay rate (in day^-1^). The model was initialized with start = list(*a* = max(volume), *b* = 0.01) for stable convergence. The fitted slope b describes the rate at which the volume decreases due to evaporation. To express this as a fractional evaporation rate per day, *b* was converted using the expression 1 − e^−*b*^, which accounts for the proportion of volume lost daily under an exponential decay assumption. This approach provided a normalized and interpretable measure of evaporation rate that could be compared across treatments and conditions.

### Macroalgal samples and cultivation

All procedures were optimized with *Laminaria ochroleuca* Bachelot de la Pylaie, 1824 female and male gametophytes from the CCMAR Biobank (CCMAR codes CCMAR-LO3.8F_CA and CCMAR-LO3.6M_CA) sampled from Cascais, Portugal in 2024, during a research cruise organized by the Blue Ocean Foundation, then transferred to the Laboratory of Biogeographical Ecology and Evolution at the Center of Marine Sciences (CCMAR) in Faro, Portugal. Spore dehiscence induction and gametophyte isolation were performed as previously described (17). Other species were used to validate the protocol: *Ulva sp and Gracilariopsis sp* (sampled in the Ria Formosa Marine Protected Area) and *Phyllariopsis purpurascens* (CCMAR-PP26_CA), sampled at artificial reefs located near the Ria Formosa lagoon, 2.0–3.0 km offshore Olhão, Portugal (63). Two other rhodophytes, a *Erythrotrichia sp* and a *Hydrolithon sp* grew as epiphytes of *Gracilariopsis* and *Ulva,* respectively, and could be cultured and isolated. *P. purpurascens* provided spores for the isolation.

Cultivation was done in filter sterilized (0.22 μm) Provasoli-Enriched Artificial Seawater (**PEAS**: ASW with half-strength Provasoli-Enrichment Solution) at salinity 28, pH 8.2, at 13°C under 8 μmol·m^-2^·s^-1^ of red light and 16:8h light:dark cycle in a climate-controlled chamber (Fitoclima S600, Aralab). The salinity was reduced as some evaporation was expected. Non-Phaeophyceae seaweeds were cultivated at 13°C under 50 μmol·m^-2^·s^-1^ of white light and 12:12h light:dark cycle. After distributing gametophyte fragments in the plates, a 15 min incubation at 13°C was included to allow random gametophyte decantation. The plates were then centrifuged (3,700 × g, 4 °C, 1 min) to eliminate residual medium and recently attached gametophytes from the well walls prior to sealing. All plate manipulation procedures were done inside a laminar flow hood to minimize contamination.

### Microscopy and Image Segmentation

#### Microscope setup

Chlorophyll auto-fluorescence (CAF) was optimized in glass-bottom black 384 well plates (384WP; VWR: 732-3746) containing 100 μl PEAS/well. The plates were manually screened in an inverted fluorescence microscope (Zeiss Observer D1) placed in a dark room at 20°C. Macroalgal tissue CAF was visualized at 5x magnification (Zeiss EC EpiPlan HD 0.13 NA), and a PI (propidium iodide) filter (Zeiss filter set 00; Ex BP 530-585 | Em LP 615). A high resolution full-frame MILC (Mirrorless Interchangeable-Lens Camera; CANON EOS-RP, 35mm sensor, 26MP) was adapted to the microscope 60N baseport by a 1.6x T2-T2 adapter (Zeiss 426115-0000-000) coupled with a T2-60N microscope adapter (Zeiss 426103-0000-000) and a M42 to RF camera thread adapter ring. To enhance the contrast between sample structures and well boundaries, exposure was set to 0.5s, inducing controlled saturation to facilitate clear delineation of edge artifacts and well contours. The wide Field of View (FOV: 3mm) allowed the coverage of a full well with a single photo. The focus was manually adjusted for each well. All plates were centrifuged (3,700 × g, 4 °C, 1 min) and had the SSMs removed before photomicroscopy.

To guide the manual screening, we used a meandric format with a “down & right” order, beginning in well A1, so the sequence of the photos could be numbered in a permanent order to associate with a plate design table containing the metadata for each well, including identification with its respective image. Images were acquired using a remote shutter and central alignment was guided by the camera’s built-in screen grid, displayed on a 23-inch monitor (ACER 8H43HX) via a HDMI connection.

#### Image processing

Acquired images were post-processed using FIJI software (ImageJ v1.54g) and segmented with *LabKit* (24,25), a built-in FIJI plugin employing a random forest pixel classification algorithm. Image processing was performed on an ASUS X99-DELUXE II workstation equipped with an Intel(R) Xeon(R) 2.20 GHz CPU, 128 GB of RAM, and a GeForce GTX 1050 Ti GPU.

To train the segmentation model on *Labkit*, gametophytes from 10 different wells were randomly selected from photos captured with a PI filter. A model was created defining two classification classes: (1) Background and (2) Live gametophytes. The Background class included dead gametophytes, surrounding background regions of live gametophytes, unidentified particles, and well edges. The Live gametophytes class encompassed verified viable tissues, confirmed by overlaying Regions of Interest (ROI) from segmented images and from bright field microscopy. This classification approach ensured that only viable tissue was segmented into the second class. After confirming the pixels relative to viable and dead gametophytes, the model could be retrained and improved. This model was used for posterior analysis.

To evaluate the precision of the Labkit segmentation, we compared the segmented areas of 39 gametophytes obtained using the PI-based model with the corresponding manually measured areas. Manual measurements were performed in FIJI using the polygon selection tool to delineate the outermost contours of each gametophyte, and the regions were saved as individual ROI files. Due to the filamentous nature of the gametophytes, these contours frequently included internal background regions formed by the natural arrangement of filaments. To account for this, these background regions were manually traced in separate ROIs and subsequently subtracted from the original measurements. This approach ensured that only the true biological tissue was quantified, allowing for a robust and accurate comparison with the LabKit-derived segmentation per gametophyte. To validate the Labkit segmentation model, automated area measurements were compared against manually obtained polygon-based ROIs in FIJI, and a linear regression analysis was used to assess the correlation between methods.

A custom FIJI macro script was developed to automatically batch-process all images. The script applied the pre-trained model to the corresponding filter-specific images, thresholded the classified outputs, conducted morphometric particle analysis, and saved all resulting data. The macro script incorporated a plate design metadata table to ensure accurate image file naming by plate, date and well position. Additional experimental data (e.g., species, sex, dilution and fragmentation technique) was also detailed in the plate design table.

### Fragmentation and sieving

To evaluate the efficiency of different gametophyte fragmentation methods for high-throughput applications, we compared three mechanical techniques: 1) TissueLyserII (TL; QIAGEN) fragmentation, 2) mortar and pestle (MP), and 3) portable motorized microtube grinder (MG; SIGMA: Z359971). Each method was applied for three fragmentation times (1, 2, and 3 minutes). Additionally, TissueLyser treatments included a frequency sub-factor with three levels (20, 25, and 30 Hz), resulting in a total of 15 unique method-time-frequency combinations. Fragmentation was performed on one 2 mm healthy (no chlorosis) gametophyte tuft. All material in contact with gametophytes, including TL racks, was pre-cooled to 4°C.

For the TL method, round bottom 2 ml microtubes (Eppendorf, EP0030120094) with 1 sterile tungsten carbide bead and 500 μl of ASW were used. After fragmentation, the beads were removed with a neodymium magnet. For the MP method, 10 cm diameter autoclaved mortars were filled with 500 μl of 4°C ASW and the tissue was gently ground and then transferred to 1.5 ml tubes with a micropipette. For the MG method, tufts were transferred to sterile 1.5 ml microtubes with 500ul of PEAS and disrupted with sterile CTFE pestles (Sigma PN Z359963).

The disrupted tissue was washed 3x with cold ASW by centrifugation (3,700xg; 4°C, 1min) to remove cell debris and eluted in 1ml of ASW with 2 μg/ml of BSA (0.2% Bovine Serum Albumin) to reduce attachment to plastic surfaces, increasing pipetting precision of low volumes. To compare the fragmentation level (counts) and recovery yields (total gametophyte area) between methods and their subsequent effect on growth, 50 μl of the fragmented tissue was plated on 384WP wells (n = 16) after sieving through 20 μm cell strainers (VWR: 734-3619) and then screened by fluorescence microscopy.

### Dilution-to-extinction isolation

To find an efficient dilution that could generate the higher quantity of wells with a single gametophyte fragment, we performed a dilution-to-extinction (DTE) isolation that consisted of serial dilutions per row starting from 2x (2^1^) to 65,536x (2^16^). We started by distributing 100 μl of the fragmented and sieved seaweed tissue into the first row of a 384WP and then, by using a 12 channel multipipette, 50 μl were removed and diluted into a previously filled well with the same volume in a lower row of the plate. This was repeated with a multichannel pipette until the last row, creating 16 dilutions (2^16^). The plates were then submitted to cultivation as described previously.

### Phenotyping by Growth rate analysis

To directly evaluate edge effects on growth kinetics within 384WP, fifteen healthy 2 mm gametophyte tufts from male and female strains were fragmented, washed, sieved using the optimized conditions described above. The resulting inoculum was diluted 10x and plated in triplicate to determine initial density and calculate the appropriate working dilution. Fragments were then suspended in a sterile reagent reservoir (VWR: 732-0794) with PEAS medium and 100 μl were distributed across all wells using a 12-channel multipipette. Plates were monitored by fluorescence microscopy every two days over 40 days after-fragmentation (DAF), capturing a complete growth curve encompassing the lag, exponential, and stationary phases. No media exchange was performed during this period.

### Data Processing and Statistical Analysis

Data distribution was assessed using the Shapiro-Wilk test to evaluate normality and homogeneity of variances was evaluated with Levene’s test. When both assumptions were satisfied (p > 0.05), group comparisons were conducted using one-way ANOVA. For non-parametric distributions, the Kruskal-Wallis test was applied for multiple group comparisons. When appropriate, Dunn’s *post hoc* with Benjamini–Hochberg correction was applied to compare multiple pairwise comparisons, i.e on fragmentation methods efficiency, edge-effects, to compare evaporation rates and growth rates.

Daily Specific Growth Rates (SGR) were calculated using R! Studio software (56) using the *fit_easylinear* function from the *growthrates* package, which fits a linear model to the exponential phase of growth curves, allowing for robust estimation of the specific growth rate (μ) based on the slope of the log-transformed data within a linear interval defined as a function parameter (64). To facilitate visualization and well-to-well comparison of raw and processed growth data, we developed a custom R! script that maps the entire 384WP layout, the *plate plot*, overlaying growth curves and calculates daily SGR and daily SERs for each individual well.

## CRediT authorship contribution statement

**C.A.L.**: Conceptualization, Methodology, Investigation, Validation, Data Curation, Software, Formal analysis, Writing – Original Draft, Visualization. **F.F.:** Methodology, Writing - Review & Editing. **LB,** Methodology, Project Administration, Writing – Review & Editing. **C.M:** Resources, Writing - Review & Editing. **LD, ML, RC, LR**: Investigation, Writing – Review & Editing. **FA, A.E.**: Investigation, Supervision, Funding acquisition, Writing – Review & Editing. **E.A.S**: Funding Acquisition, Sampling, Writing – Review & Editing. **G.P., N.M.**: Investigation, Supervision, Methodology, Funding Acquisition, Supervision, Writing – Review & Editing.

## Competing interests

The authors declare that they have no known competing financial interests or personal relationships that could have appeared to influence the work reported in this paper.

## Acknowledgments

This study received Portuguese national funds from FCT - Foundation for Science and Technology through contracts UID/04326/2025, UID/PRR/04326/2025 and LA/P/0101/2020 (DOI:10.54499/LA/P/0101/2020), and AdaptKelp (MPr-2023-12-16551, ALGARVE-FEDER-00772500), and BIODIVERSA BiodivRestore-253 - FCT DivRestore/0013/2020. **CAL** salary and the CCMAR Biobank were financed by the WP9 - Portuguese Blue Biobank under the Blue Economy Pact - Project N° C644915664-00000026 co-funded by PRR, the Portuguese Republic and the European Union. We thank Dr. Carla Cherubini for her valuable assistance on image analysis. We thank Vera Gomes and Marta Bernardo for the assistance with the CCMAR facilities for spectrophotometry and DNA sequencing. The Oceano Azul Foundation for supporting the field sampling of *Laminaria ochroleuca* in Cascais and *Halymenia sp* in the Gorringe Expeditions.

## Abbreviations

SAMMBA: Seaweed Automatable Microplate Microscopy for Breeding Approaches
384WP: 384-well plate

## Notes

### Competing Interest Statement

The authors have declared no competing interest.

### Summary of Updates

Major reviews on abstract, main text, figures, tables and supplementary material. Also added Brigitta Varga and Joana Filipe for their contributions on the protocol enhancement.

