## Supplementary Figures for "SAMMBA is a high-throughput pipeline for isolating and phenotyping macroalgal strains"

Affiliations:

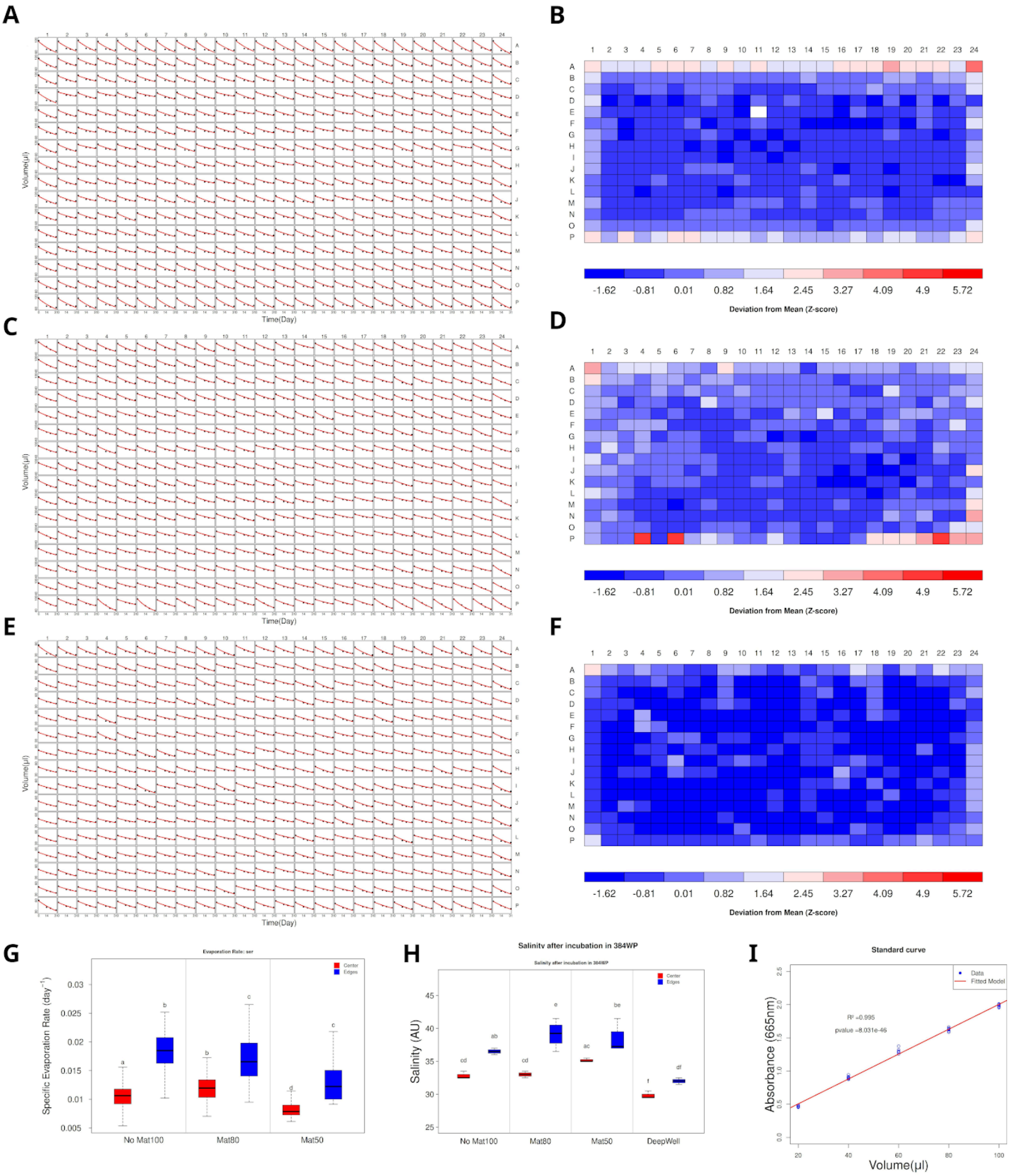


**Supplementary Figure 1. Reduction of edge effects by silicone sealing mats in 384-well plates.** Panels A–B represent an uncovered plate filled with 100 µL per well; C–D a sealed plate filled with 80 µL; E–F a sealed plate filled with 50 µl. In A, C and E, evaporation curves showing volume loss over time per well, estimated using the methylene blue standard curve in panel G. In B, D and F, Z-scores of specific evaporation rates (SER) per well derived from the exponential decay slopes of evaporation curves. In G, comparison of SER from edges wells (rows A and P and columns 1 and 24) versus center wells per treatment. In H, final salinity measurements compare only edge wells (A1, A24, P1, P24) and center wells (H12, H13, I12, I13) across all treatments. In I, standard curve relating methylene blue absorbance at 665 nm to volume in 384-well plates.

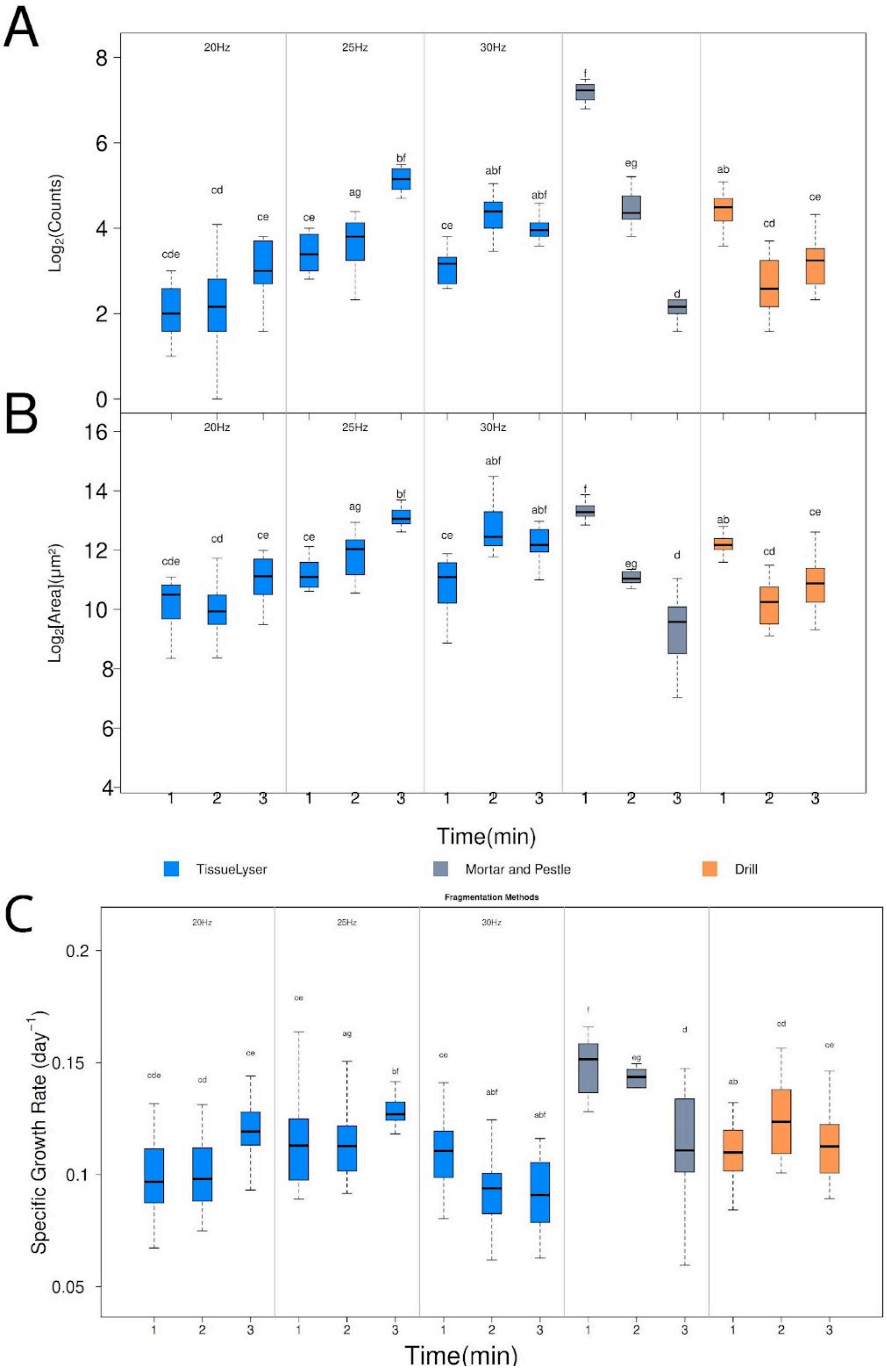


**Supplementary Figure 2. Comparison of fragmentation methods of *Laminaria ochroleuca* gametophytes**. A: Total fragment counts per well. B: Total recovered area per well. C: Daily Specific Growth Rate (SGR). Compact Letter Displays (CLD) above boxplot whiskers indicate significant pairwise differences based on Dunn's test (adjusted p-values). For the TissueLyser method, different frequencies tested are highlighted within each panel. Method color codes are shown in the legend below the plots.

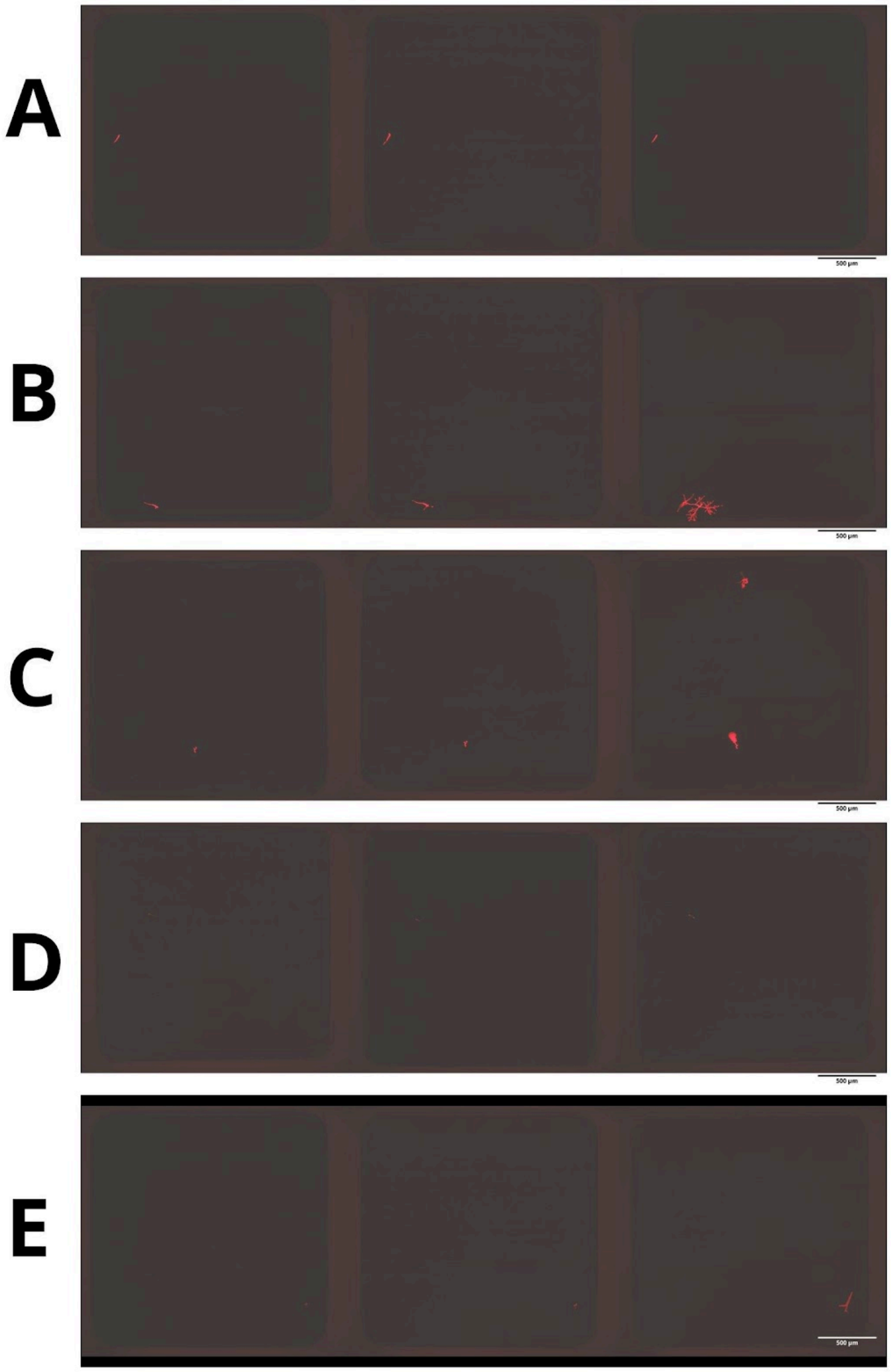


**Supplementary Figure 3. Examples of isolates monitoring of non-Laminariales seaweed along one month.** First panels show the first day of isolation, second panel 1 week later and last panel one month later. A: *Erythrothrychia sp*. B: *Halymenia sp*. C: *Hydrolithon sp*. D: *Phyllariopsis purpurascens*. E: *Ulva lactuca.* Note that on the third panel of C, a second isolate is visible, changing the characterization of the isolate from singlet to doublet.

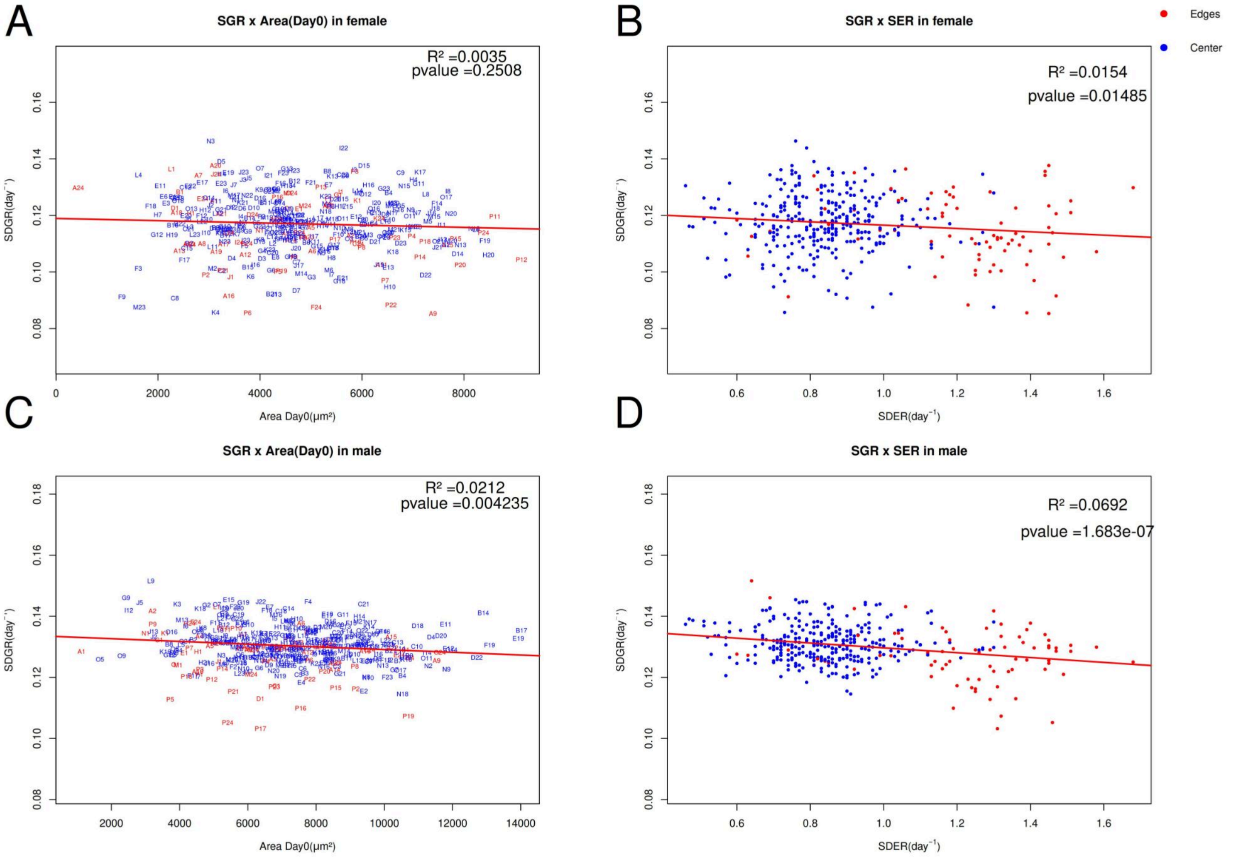


**Supplementary Figure 4**. No correlation was observed between daily Specific Daily Growth Rate (SGR) and initial area and specific daily specific evaporation rate (SER).

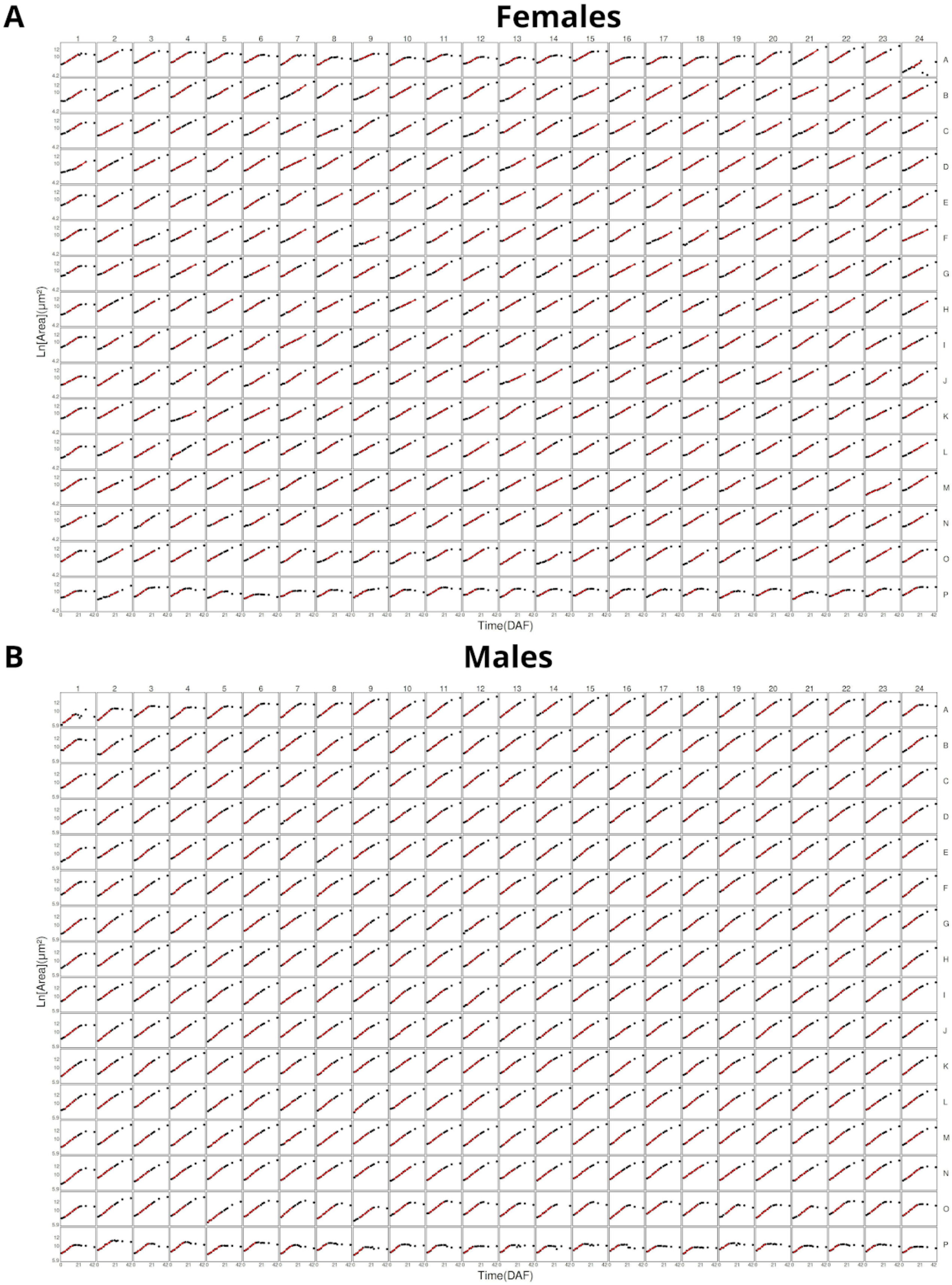


**Supplementary Figure 5. Growth curves of Laminaria ochroleuca gametophytes over 42 days following fragmentation.** (A) Female and (B) male gametophytes, showing earlier onset of stationary phases in wells located at the plate edges. No aberrant time points were detected, underscoring the accuracy of total gametophyte area measurements. Red lines indicate the time interval used for growth rate calculations. DAF stands for “days after fragmentation”.

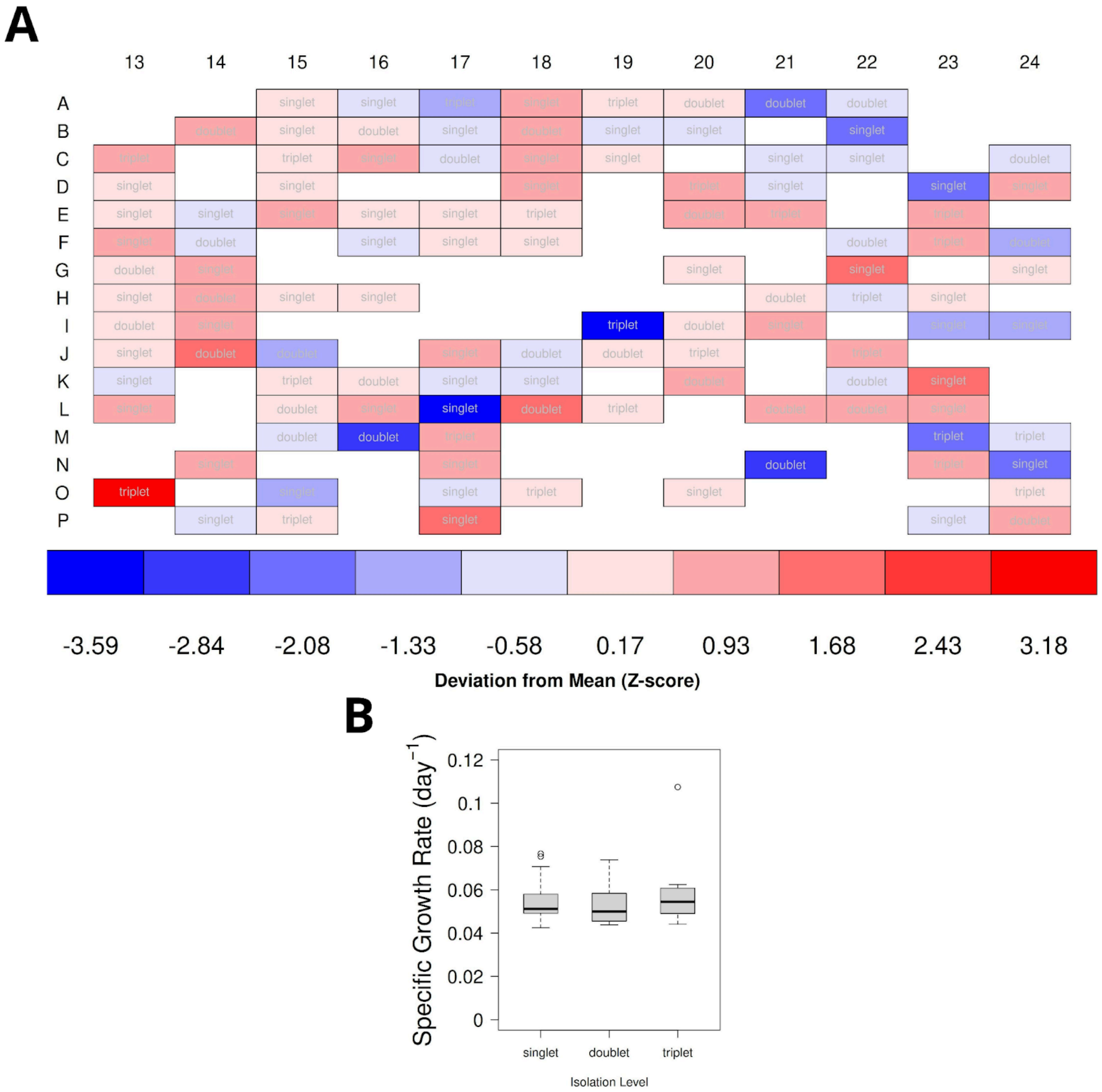


**Supplementary Figure 6. Overview of the daily Specific Growth Rates (SGR) from the isolates of *Phyllariopsis purpurascens*.** In A, a Plate-plot showing spatial distribution of SGR for singlets, doublets and triplets isolates across half 384 well plate. Z-scores were computed for each well and visualized using red–blue palettes to reflect deviations from the mean SGR. The white squares mean that the correspondent well was empty or had more than 4 fragments.

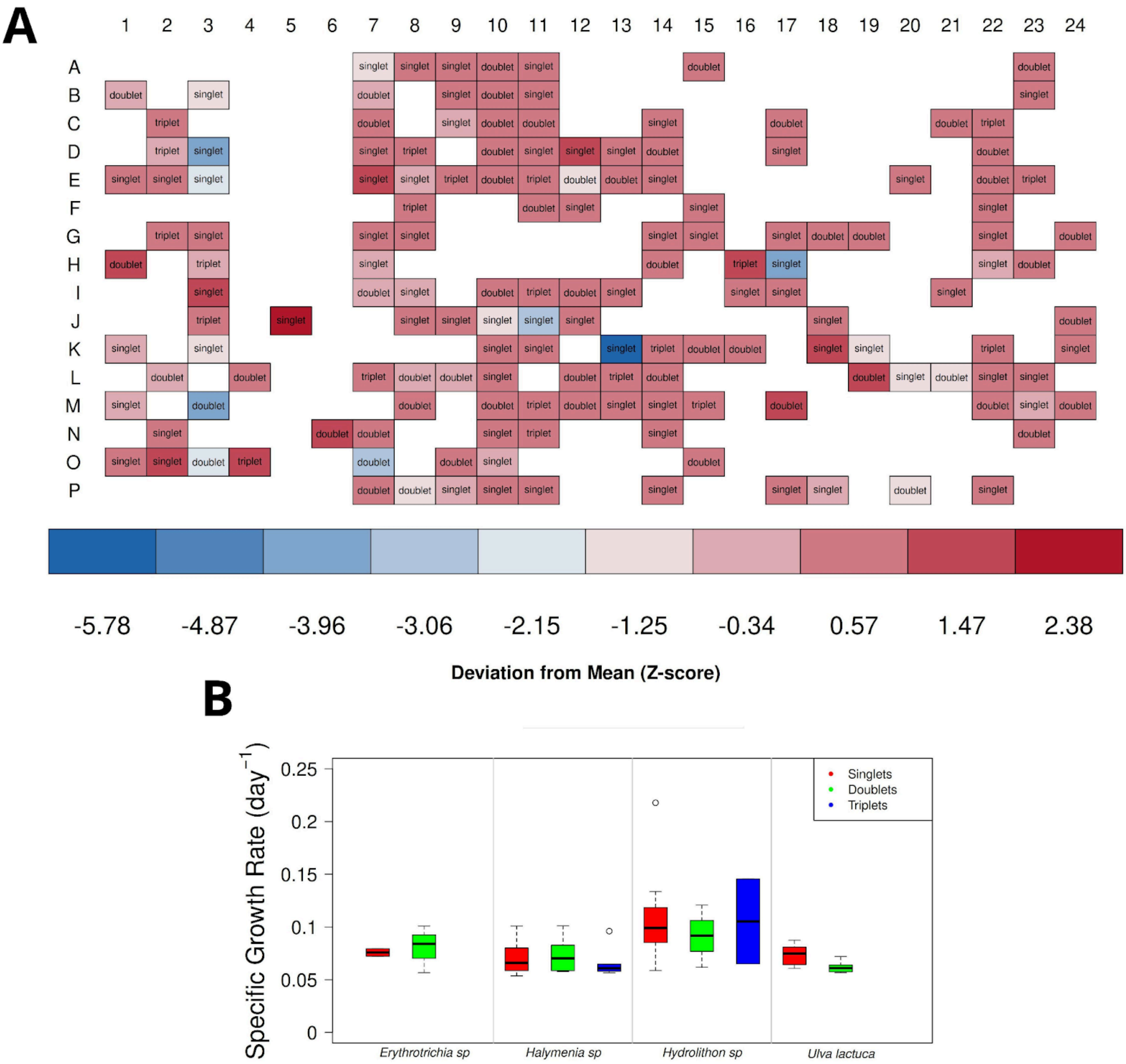


**Supplementary Figure 7. Overview of the daily Specific Growth Rates (SGR) from the isolates of Rhodophyte and Ulvophyceae seaweed.** In A, a Plate-plot showing spatial distribution of SGR for singlets, doublets and triplets isolates across the plate. Z-scores were computed for each well and visualized using blue–red palettes to reflect deviations from the mean SGR. The white squares mean that the correspondent well was empty or had more than 4 fragments.

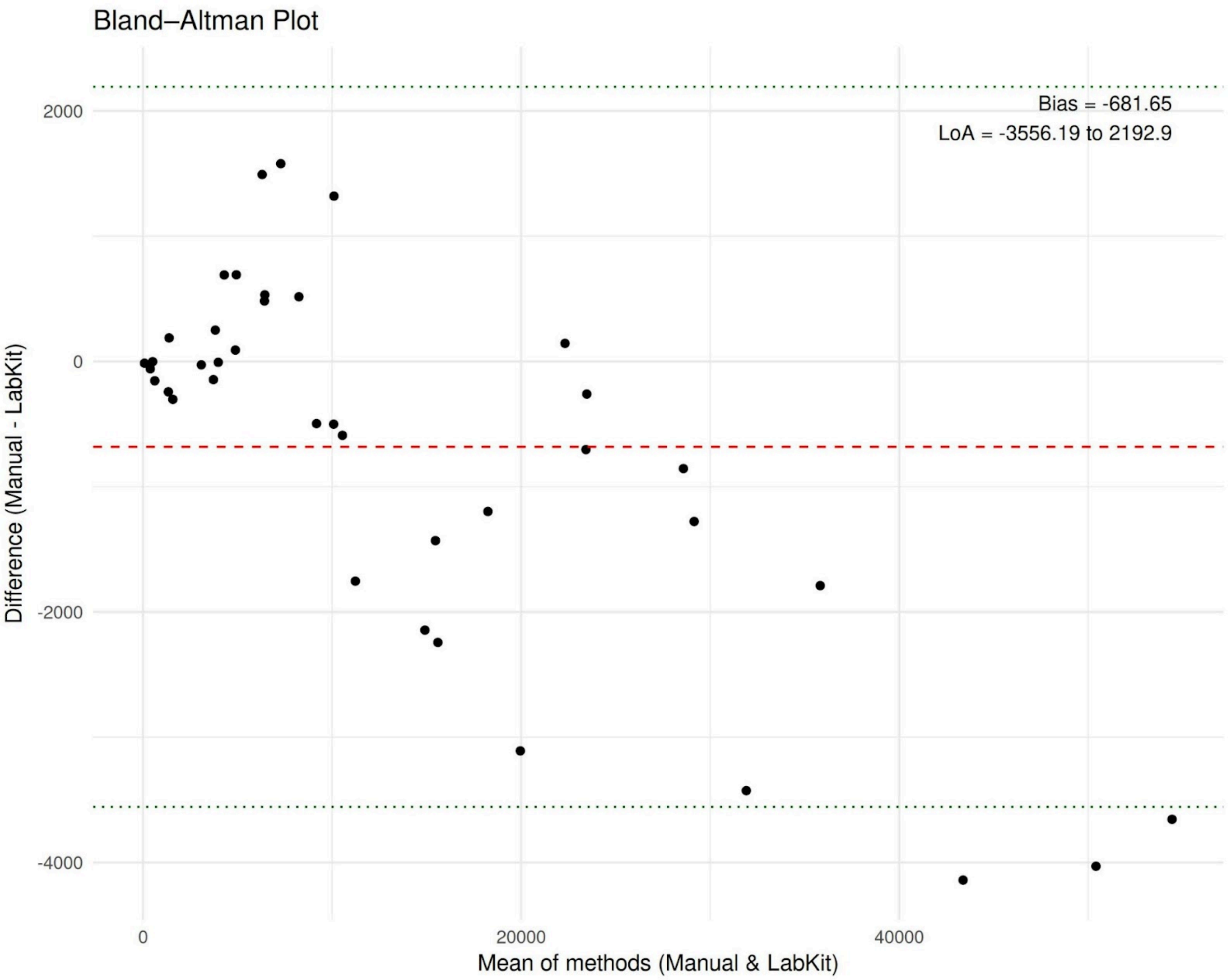


Figure 8. Bland–Altman analysis of measurements from the two methods, showing a bias of –681.65 and limits of agreement ranging from –3556 to 2192.9.
